# Organ-specific propensity drives patterns of gene expression evolution

**DOI:** 10.1101/409888

**Authors:** Kenji Fukushima, David D. Pollock

## Abstract

The origins of multicellular physiology are tied to evolution of gene expression. Genes can shift expression as organisms evolve, but how ancestral expression influences altered descendant expression is not well understood. To examine this, we amalgamated 1,903 RNA-seq datasets from 182 research projects, including 6 organs in 21 vertebrate species. Quality control eliminated project-specific biases, and expression shifts were reconstructed using gene-family-wise phylogenetic Ornstein–Uhlenbeck models. Expression shifts following gene duplication result in more drastic changes in expression properties than shifts without gene duplication. The expression properties were tightly coupled with protein evolutionary rate, depending on whether and how gene duplication occurred. Fluxes in expression patterns among organs were nonrandom, forming modular connections which were reshaped by gene duplication. Thus, if expression shifted, ancestral expression in some organs induces a strong propensity for expression in particular organs in descendants. This supports a major role for what might be termed “preadaptive” pathways of gene expression evolution.

## Introduction

Vertebrate organs organize physiological activities, and their identities and functions are determined by the diverse expression patterns of thousands of genes. Because of this, the evolution of gene expression patterns plays a central role in organismal evolution. The degree of organ expression specificity correlates to how fast amino acids substitute (Zhang and Li, 2004), how rapidly they change expression levels (Liao and Zhang, 2006), and patterns of histone modifications (She et al., 2009). Major organ-altering evolutionary events such as development of the hominoid brain are also associated with gene expression shifts (Chen et al., 2013; Kaessmann, 2010; Zhang and Long, 2014; Zhang et al., 2011). However, although gene duplication is well-known to play an important role in expression pattern shifts (see e.g. the “ortholog conjecture” (Altenhoff et al., 2012; Castillo-Davis et al., 2004; Chen and Zhang, 2012; Rogozin et al., 2014)), the evolutionary dynamics of expression patterns with and without gene duplication remain poorly understood. An important question is whether long-term expression in one organ predisposes genes to be subsequently utilized in other organs.

A theoretical basis for such predisposition may be the idea that certain pre-existing adapted states are more conducive to evolution of specific new traits than other pre-existing states. This is known as preadaptation, and when a trait makes such a shift it is referred to as exaptation (Gould and Vrba, 1982). Evidence for preadaptation was long ago found in phenotypic traits (Budd, 2007), and recently in molecular traits such as protein sequences during *de novo* gene birth (Wilson et al., 2017) or during functional innovations (Starr et al., 2017). Protein sequence evolution generally involves highly epistatic interactions and context-dependent changes (Podgornaia and Laub, 2015; Starr et al., 2017) that affect preadaptation, but the modular nature of expression regulation (de-Leon and Davidson, 2007) makes it unclear whether preexisting expression patterns constrain evolutionary outcomes. By analyzing such influences on a genome-wide scale in this study, we aim to contribute some evidence towards this question.

Evolution of gene expression has been studied at genome-wide scales mainly using two distinct approaches: phylogenetic and pairwise analyses. Phylogenetic approaches model gene expression dynamics and infer ancestral expression patterns in the context of gene phylogenies. For example, Brownian motion models embody purely neutral expression evolution (Bedford and Hartl, 2009), whereas Ornstein–Uhlenbeck models are designed to detect purifying selection and adaptive evolution along with neutral fluctuation (Brawand et al., 2011; Chen et al., 2019; Rohlfs et al., 2014). Although each gene family has a distinct evolutionary history, a species phylogeny is often used for the sake of simplicity. Because such approximations cannot be applied to gene families with lineage-specific gene duplications and losses, its application has mostly been limited to single-copy genes. In contrast, pairwise analysis compares gene expression between paralogs in single species (Assis and Bachtrog, 2015; Lan and Pritchard, 2016) or between orthologs or paralogs in pairs of species (Chen and Zhang, 2012; Guschanski et al., 2017; Kryuchkova-Mostacci et al., 2016; Rogozin et al., 2014; Warnefors and Kaessmann, 2013). Although pairwise approaches can evaluate the effect of gene duplications, ancestral expression cannot be inferred.

To infer the adaptive evolution of gene expression in diverse gene families, we applied Ornstein–Uhlenbeck models for complex gene family phylogenies containing gene duplications and losses, without assuming species phylogeny. We also developed a curation pipeline to amalgamate large amounts of transcriptome data from many studies for a better phylogenetic resolution. Using these methods, we ask how expression changes link to gene duplication and the rate of protein evolution, and whether previous patterns of gene expression constrain later evolution, what we will call the “preadaptation hypothesis”.

## Results

### Duplication-permissive genome-wide analysis of gene families

To allow evolutionary expression analysis on a broad set of genes, we used a phylogenetic approach that deals with the complex history of gene family trees with duplications and losses, and applied it to 21 tetrapod genomes (Fig. S1). A major challenge in using gene trees was divergence time estimation, a prerequisite for applying phylogenetic comparative methods. We overcame this problem by incorporating phylogeny reconciliation in estimating divergence time of gene trees. Gene divergence nodes were constrained by the corresponding divergence times in a known species tree, and duplication nodes were constrained by ancestral and descendant speciation events (see Materials and Methods for details). Because we estimated individual gene phylogenies rather than using a single species phylogeny, we could analyze gene families that included many lineage-specific gene duplications and losses, making our study less biased towards conserved genes with slow gene turnovers (Guschanski et al., 2017). Use of gene family trees also allowed us to include many organ-specific genes that are enriched in lineage-specific and young duplications (Guschanski et al., 2017). There were only 1,377 single-copy orthologs for which the species phylogeny was applicable, but we were able to include 15,475 genes per species on average (including 20,873 human genes, merged into 15,280 gene families). This approach eliminates problems with pairwise analyses that ignore phylogenetic tree structure, and allowed us to infer expression at ancestral nodes in the tree.

### Transcriptome amalgamation

To attain high resolution in our analyses, we amalgamated 1,903 RNA-seq experiments from 182 research projects (i.e., 182 BioProject IDs in the NCBI SRA database) and generated a dataset covering 6 organs from 21 vertebrate species without missing data (Tables S1–S2; Fig. S1). In comparison, other recent comparative transcriptomic analyses of vertebrates (Barbosa-Morais et al., 2012; Breschi et al., 2016; Carelli et al., 2016; Chen et al., 2019; Cortez et al., 2014; Guschanski et al., 2017; Julien et al., 2012; Necsulea et al., 2014; Warnefors and Kaessmann, 2013) often used the same dataset containing 131 RNA-seq experiments from 6 organs and 10 species (Brawand et al., 2011), with some additional data in different studies. RNA-seq reads were first mapped to corresponding reference genomes and then the expression level was quantified by two metrics: transcripts per million (TPM) and fragments per kilobase million normalized by trimmed mean of M-values (Robinson and Oshlack, 2010) (TMM-FPKM). To reduce the among-species variation, the TMM normalization was applied across all 1,903 samples using the 1,377 single-copy orthologs.

To allow rapid integrated analysis of datasets, we employed automated multi-aspect quality controls, including metadata curation (Table S3), sequence read filtering (Fig. S2), and iterative removal of anomalous RNA-seq samples by monitoring correlations between and within data categories (Fig. 1A; Fig. S3). The metadata curation enabled us to select appropriate samples from the NCBI SRA database. Data that were not compatible with those from other research projects (defined by BioProject ID) were removed in the correlation analysis by implementing a majority rule (Table S3), resulting in a cleaned dataset. This filtering step was designed to fulfill the assumption that any samples from the same organ should correlate better than samples from different organs within species. When anomalous data were detected, all samples belonging to the same research projects (i.e., the same BioProject ID) were discarded.

**Figure 1.**
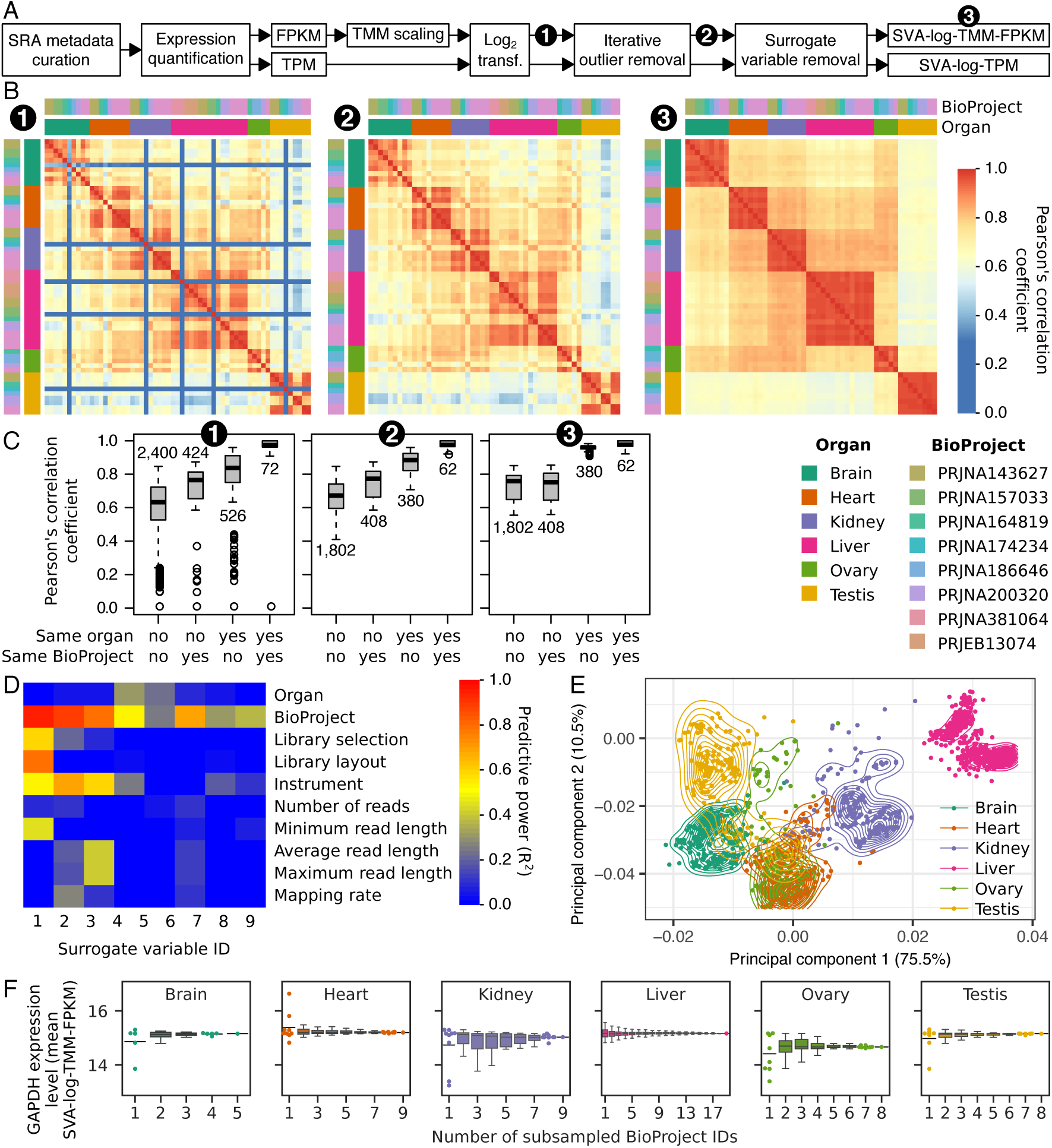
Transcriptome amalgamation to integrate heterogeneous RNA-seq samples. (**A**) A simplified flow chart of the transcriptome amalgamation. The full chart is available in Fig. S1. (**B–D**) Transcriptome curation within species. Data from *Monodelphis domestica* with SVA-log-TMM-FPKM metrics are shown as an example. The heatmaps show Pearson’s correlation coefficients among RNA-seq samples (**B**). Each row and column correspond to one RNA-seq sample. The expression levels of all genes were used to calculate the correlation coefficients. Note that anomalous samples contaminated in the curated metadata (low correlation samples in 1) are successfully removed, and that project-specific correlations visible in the uncorrected data (marked 2) are absent in the corrected data (marked 3). The box plots show distinct distributions of the correlation coefficients depending on whether a pair of samples are the same organ or whether they are from the same research project (**C**). The numbers of comparisons are provided in the plot. The correlation coefficients are largely improved in within-organ comparisons when surrogate variables are removed, while within-project biases are attenuated. In this species, nine surrogate variables were detected against 52 RNA-seq data from eight projects (**D**). Analysis of those variables by linear regression highlights the BioProject feature as the strongest source of removed biases. For full description of predictors, see Fig. S4. (**E**) A principal component analysis using expression levels of 1,377 single-copy orthologs from 21 species. Points correspond to RNA-seq samples. Curves show the estimated kernel density. Explained variations in percentages are indicated in each axis. (**F**) Estimated organ-wise expression levels of a housekeeping gene. Since data from relatively many BioProjects are available, glyceraldehyde-3-phosphate dehydrogenase gene (GAPDH, Ensembl gene ID: ENSGALG00000014442) in *Gallus gallus* is shown as an example. Points corresponds to the average expression level calculated by random subsampling. All data points and the median value (bar), rather than a box plot, are shown if the number of subsampled BioProject combinations is less than 10.

Finally, we applied surrogate variable analysis (SVA) (Leek and Storey, 2007) to detect and correct hidden biases likely originating from heterogeneous sampling conditions and sequencing procedures among experiments in both log_2_-scaled TPM and TMM-FPKM (SVA-log-TPM and SVA-log-TMM-FPKM, respectively; Fig. S4A–B). This correction greatly improved the correlation of expression levels within organs from the same species, even when data were derived from different research projects (Fig. 1B–C; Fig. S4C–F). Among surrogate variables, BioProject IDs tend to show a high predictive power, suggesting project-specific sources of bias (Fig. 1D; Fig. S4G–H). Although the inclusion of many species from phylogenetically diverse lineages makes it difficult to extract organ-wise characteristics from the limited number of single-copy orthologs, a principal component analysis produced moderate organ-wise segregation in the multispecies comparison (Fig. 1E; Fig. S4I–K), further indicating that the curated dataset is sufficiently reliable for use in cross-species expression pattern analyses. The previously-reported uniqueness of testis transcriptomes (Brawand et al., 2011) was partly resolved as the third principal component (Fig. S4K).

To further evaluate the validity of amalgamated transcriptomes, we analyzed the expression of community-curated cell-type-specific marker genes associated with organs in PanglaoDB (Franzén et al., 2019), which organizes a number of single-cell RNA-seq experiments in human and mouse. We compared median values of log-transformed expression levels of >100 marker genes in each organ (Fig. S5). After SVA correction, all RNA-seq samples in the both species showed the corresponding marker expression values higher than those from the other organs, suggesting our amalgamated transcriptomes preserve the organ-specific gene expression. In the cell-type-wise analysis, a few cases, such as juxtaglomerular cells in kidney and hepatic stellate cells in liver, could not resolve our organ-wise transcriptomes (Supplementary Data). However, such low performance was seen in all samples rather than subsets associated with particular BioProject IDs, suggesting that the dissection decisions have negligible effects to cell type compositions in the organs.

In addition to the better phylogenetic coverage, greater accuracy of estimated expression levels is another possible advantage of integrating many RNA-seq datasets. This idea is supported by subsampling analysis on a housekeeping gene GAPDH (glyceraldehyde-3-phosphate dehydrogenase), where, as more data are used, estimated expression levels tend to quickly converge to a final value (Fig. 1F; Fig. S4L).

### Modeling expression evolution

We next used the amalgamated transcriptomes to evaluate how expression evolved along 15,280 maximum-likelihood gene family phylogenies (Table S4), employing multi-optima Ornstein–Uhlenbeck (OU) models (Hansen, 1997) to allow for possible adaptive shifts of optimal expression levels and neutral fluctuations (Brawand et al., 2011; Chen et al., 2019; Rohlfs et al., 2014). This modeling identified statistically supported expression ‘regime shifts’ (Hansen, 1997; Khabbazian et al., 2016) on each gene tree (Fig. 2A), which were then analyzed in the context of preceding duplication events. ‘Speciation’ events (S node; Fig. 2B) with no duplication were considered the ‘baseline’ mode of expression evolution because regulatory environments and expression patterns are more preserved among orthologous genes in comparison with paralogous genes produced by gene duplication (Altenhoff et al., 2012; Castillo-Davis et al., 2004; Chen and Zhang, 2012; Rogozin et al., 2014). Because OU shift detection has been applied for gene expression by assuming species tree phylogeny in single-copy genes, shifts in S branches are equivalent to those characterized previously (Brawand et al., 2011; Chen et al., 2019) but also includes many more speciation events in duplication-prone gene families. Gene tree nodes associated with preceding duplication events were categorized as DNA-based duplication or retrotransposition (D or R nodes, respectively) depending on complete intron losses (Fig. 2B).

**Figure 2.**
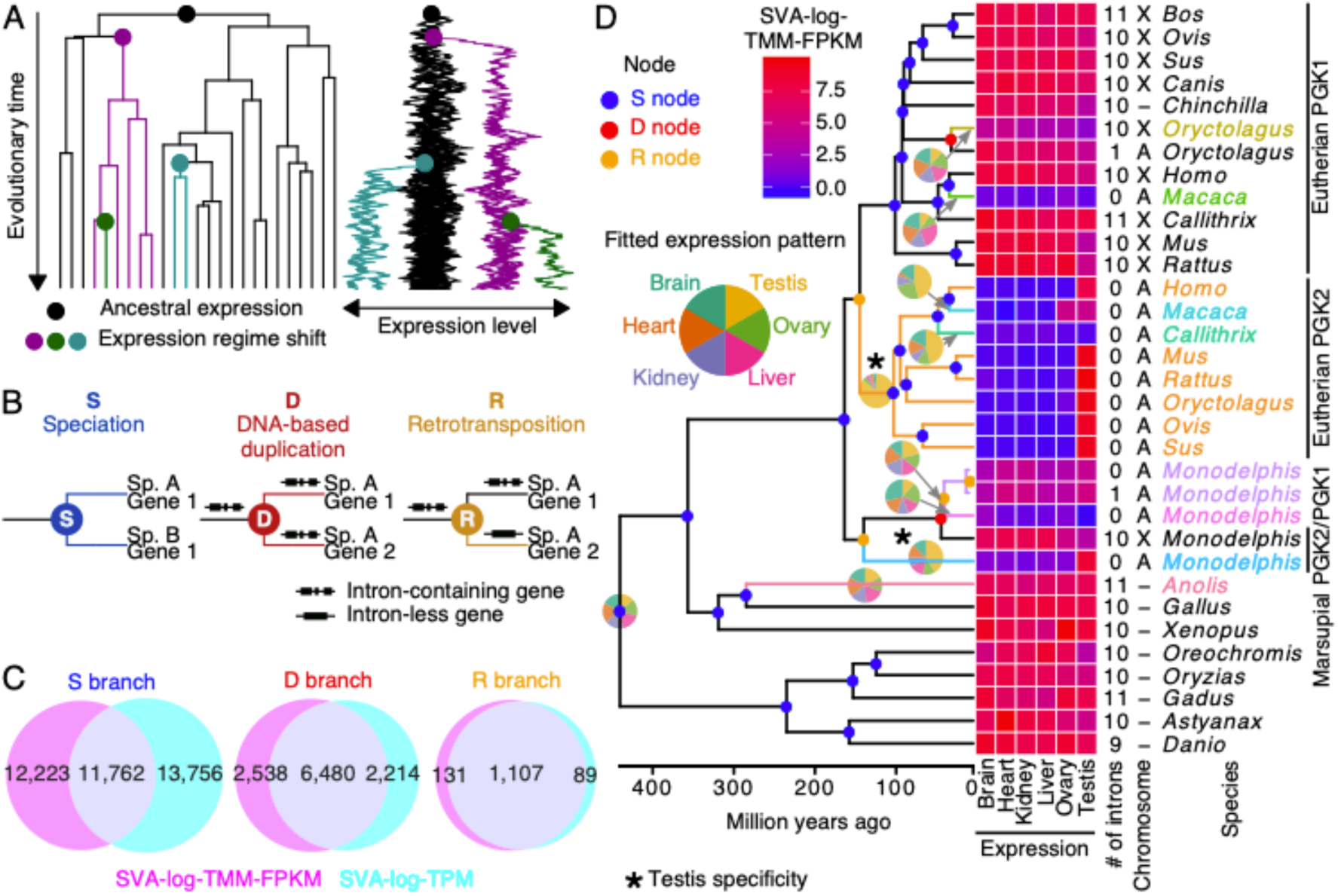
Expression evolution in a complex history of gene family evolution. (**A**) Modeling expression evolution with multi-optima Ornstein–Uhlenbeck process. A phylogenetic simulation is shown. Colors show branches belonging to different regimes. Regime shifts (change of color) appear as a substantial change in optimal trait values. (**B**) Nodes and branches of gene family phylogeny were categorized into S, D, or R based on the branching events, i.e., speciation, DNA-based duplication, or retrotransposition, respectively. (**C**) Venn diagrams of expression regime shifts. Circles represent the sets of branches where regime shifts were detected with SVA-log-TMM-FPKM or SVA-log-TPM. (**D**) The gene tree of phosphoglycerol kinases (orthogroup ID: OG0002332) is shown as an example. This orthogroup shows ortholog-specific expression patterns as well as regime shifts after speciation and lineage-specific gene duplication. Tips correspond to genes. The colors of branches and tip labels indicate expression regimes. Node colors match to the categorization in **B**. The heatmap shows expression levels and among-organ expression patterns are shown as a pie chart for each regime. To the right, the number of introns and located chromosomes (A, autosome; X, X chromosome; Y, Y chromosome) are indicated. For full information including complete tip labels and bootstrap branch supports, see Supplementary Data.

Organ-wise means of the two expression values, SVA-log-TPM and SVA-log-TMM-FPKM, were separately used to model expression evolution with OU processes. The two analyses resulted in similar numbers and characteristics of expression regime shifts (Fig. S6), but the shift locations were sometimes inconsistent. S branches were a major source of apparently inconsistent regime shifts, whereas branches following duplication (D and R) showed largely consistent detection between the two metrics (Fig. 2C). Although the inclusion of inconsistently detected branches did not change the results, we retained only consistently detected regime shifts for all downstream analysis to draw a more robust conclusion. While SVA-log-TMM-FPKM values were reported in the main text unless otherwise mentioned, the comparisons with SVA-log-TPM-based analyses are available as Supplementary Information.

As an example of this analysis, an orthogroup of phosphoglycerol kinases (PGKs), containing all three categories of branching events followed by expression shifts (S, D, and R), is shown in Fig. 2D. This protein catalyzes the first ATP-generating step in the glycolytic pathway and is required for most cell types including sperms (Danshina et al., 2010; Liu et al., 2015). PGK1, the original copy on the X chromosome, is known to have duplicated independently in eutherians and marsupials to produce the autosomal retrocopy PGK2 that compensates the protein activity during X-inactivation (Boer et al., 1987; McCarrey and Thomas, 1987; Potrzebowski et al., 2008). Our automated analysis correctly recovered both retrotranspositions as well as the subsequent gains of testis-specific expression in eutherian and marsupial lineages. This illustrates that our automated genome-wide analysis can recover evolutionary trajectories that are compatible with focused single gene family analyses (see Supplementary Data for individual gene trees).

### Duplication-specific effects in expression evolution

Across gene trees, expression regime shifts were relatively rare in S branches, for a probability of 2.2% per branch, and an average rate of 5.1×10^-4^ shifts/MY (million years) (Fig. 3A–B). In agreement with the idea that gene duplication tends to free genes up for functional divergence and enhance long-term retention of duplicated copies (Assis and Bachtrog, 2015; Huerta-Cepas et al., 2011), the frequency of regime shifts in D branches was four times as much per branch (9.0% per branch, at a rate of 1.9×10^-3^ shifts/MY across all genes and all D branches). Thus, although far fewer branches are preceded by DNA-based duplication events (72,476 branches) than speciation events (556,240 branches), D branches account for over 33% of all regime shifts consistently detected by the two expression measures. We note that this result reinforces previous results on the “ortholog conjecture”, the idea that duplicated gene copies (paralogs) are more prone to expression shifts than orthologs (Altenhoff et al., 2012; Castillo-Davis et al., 2004; Chen and Zhang, 2012; Rogozin et al., 2014). R branches were far more likely to result in expression regime shifts (37.3% per branch, at a rate of 7.6×10^-3^ shifts/MY), but with only 2,985 R branches, this resulted in only 1,107 shifts (5.6% of the total). Translocated genes are more likely to shift expression than those that do not (Fig. S7), in line with previous observations from the human genome (Lan and Pritchard, 2016). While the expression shift frequency in S and D branches varies slightly across the phylogeny, R branches showed much stronger among-lineage heterogeneity, and had a particularly high frequency in the mammalian lineage (Fig. 3B). These retrotransposition-related expression changes may be related to the variation in the retrotransposition rate itself, which is known to vary across lineages (Marques et al., 2005; Yu et al., 2007). Among-species heterogeneity in gene prediction quality may also be attributed to this pattern because early-diverging species tended to show higher percentages of missing single-copy orthologs than those in mammalian species (Fig. S8; but see *Danio rerio*, *Oreochromis niloticus*, and *Oryctolagus cuniculus* as counterexamples). In the absence of regime shifts, expression levels varied most in ovary and testes, which had significantly higher average stationary variances than the other four organs on the basis of tree-wise stationary variance (Fig. 3C). This supports previously observed high variation of gene expression in testes (Brawand et al., 2011) and extends it to the reproductive organs of both sexes.

**Figure 3.**
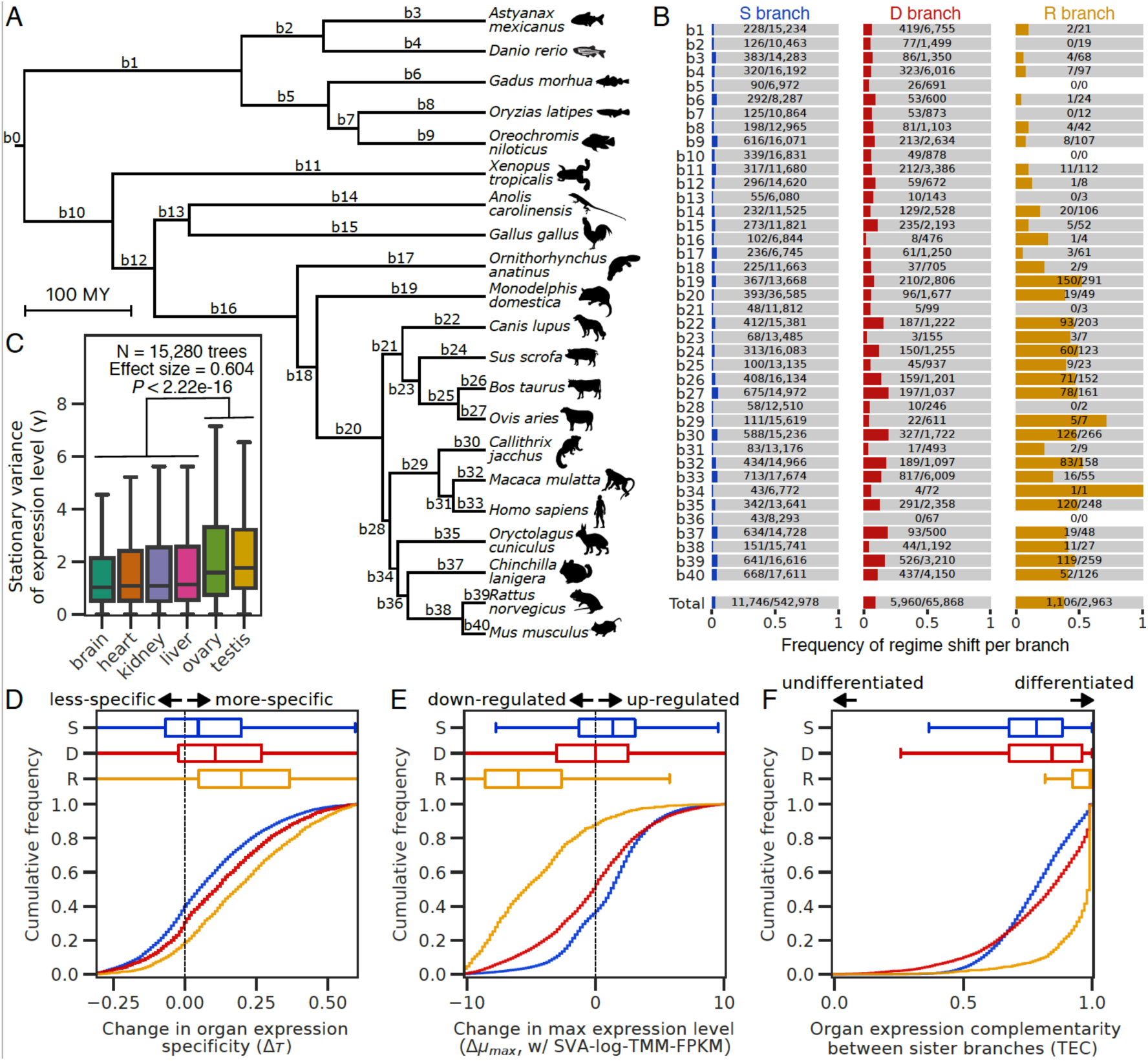
Characteristics of expression shifts in 15,280 gene trees. (**A**) The species tree showing analyzed genomes and their divergence time. (**B**) Mapping of 18,812 expression shifts in the species tree. The number and proportion of expression regime shifts in S, D, and R branches are shown. Corresponding branches in the species tree are indicated in **A**. (**C**) Organ-specific stationary variances (γ) of expression level evolution in vertebrates. The distribution of γ between reproductive and non-reproductive organs were compared by a Brunner-Munzel test (Brunner and Munzel, 2000). (**D–F**) Cumulative frequency of change in organ expression specificity (**D**), change in maximum expression level (**E**), and expression complementarity between sister lineages (**F**) among detected expression shifts. Number of analyzed regime shifts are shown in **B**.

If expression regime shifts are due to functional divergence, it is highly relevant to characterize how expression properties changed from ancestral to derived regimes. To do this, we examined changes in organ expression specificity, maximum expression level, and organ expression complementarity. The specificity measure τ ranges from 0 for uniformly expressed genes to 1 for genes with highly specific expression (Yanai et al., 2005). The distributions of regime shifts in D and R branches are shifted towards greater organ specificity compared to shifts in S branches, with R branches creating the most specific expression (Fig. 3D). To characterize the ‘on state’ transcriptional activity, we analyzed the maximum fitted expression levels among the six organs (μ_max_). D and R branches appear to be enriched for down-regulation compared to S branches (Fig. 3E). Complementarity of organ expression patterns was measured to evaluate the differentiation between a pair of sister branches. We used a metric on the fitted organ-wise expression levels (μ) called ‘TEC’, which ranges from 0 for completely overlapping expression to 1 for mutually exclusive patterns (Huerta-Cepas et al., 2011). Nearly all branches with regime shifts (95%) had complementarity values greater than 0.5, indicating that most regime shifts detected involve differentiation of expression patterns, rather than overall up- or down-regulation. Regime shifts in D and R branches often had more complementarity expression than those in S branches (Fig. 3F), further supporting the role of gene duplication in functional differentiation. The more drastic effect in R branches probably reflects the regular loss of regulatory elements in retrotranspositions, whereas DNA-based duplication can more often retain regulatory regions (Babushok et al., 2007). Jointly, these results indicate that, compared with the baseline from speciation-associated shifts, gene duplication tend to produce more organ-specific, more often down-regulated, and more differentiated expression patterns. Although the down regulation may be explainable by a tendency to need less of the newly functional expression regime, it may also be explained by either recent nonfunctionalization (Balakirev and Ayala, 2003; Mighell et al., 2000) or specialized expression in organs that were not part of this analysis.

### Context-dependent change in the rate of protein evolution

Change in gene expression can be accompanied by accelerated or decelerated protein evolution, which may be detected by change in the ratio of nonsynonymous/synonymous substitutions (*dN/dS* or ω) along branches. In D and R branches, median ω values are more than double the baseline seen in S branches (Fig. 4A; Fig. S9A), again supporting the ortholog conjecture and the idea that gene duplication tends to free genes up for functional divergence (Assis and Bachtrog, 2015; Huerta-Cepas et al., 2011). Within each of S, D, and R branch categories, branches with expression regime shifts accompany an increased ω compared to sister branches (Fig. 4A). The increased rate was quite small in S branches (median ω, 0.096 in branches with shifts versus 0.102 in sister branches), much bigger in D branches (0.182 versus 0.244), and huge in R branches (0.036 versus 0.394). If the changes in expression and the rate of protein evolution are due to functional changes, this may indicate that functional divergence is sometimes effected by joint changes in expression and accelerated protein evolution.

**Figure 4.**
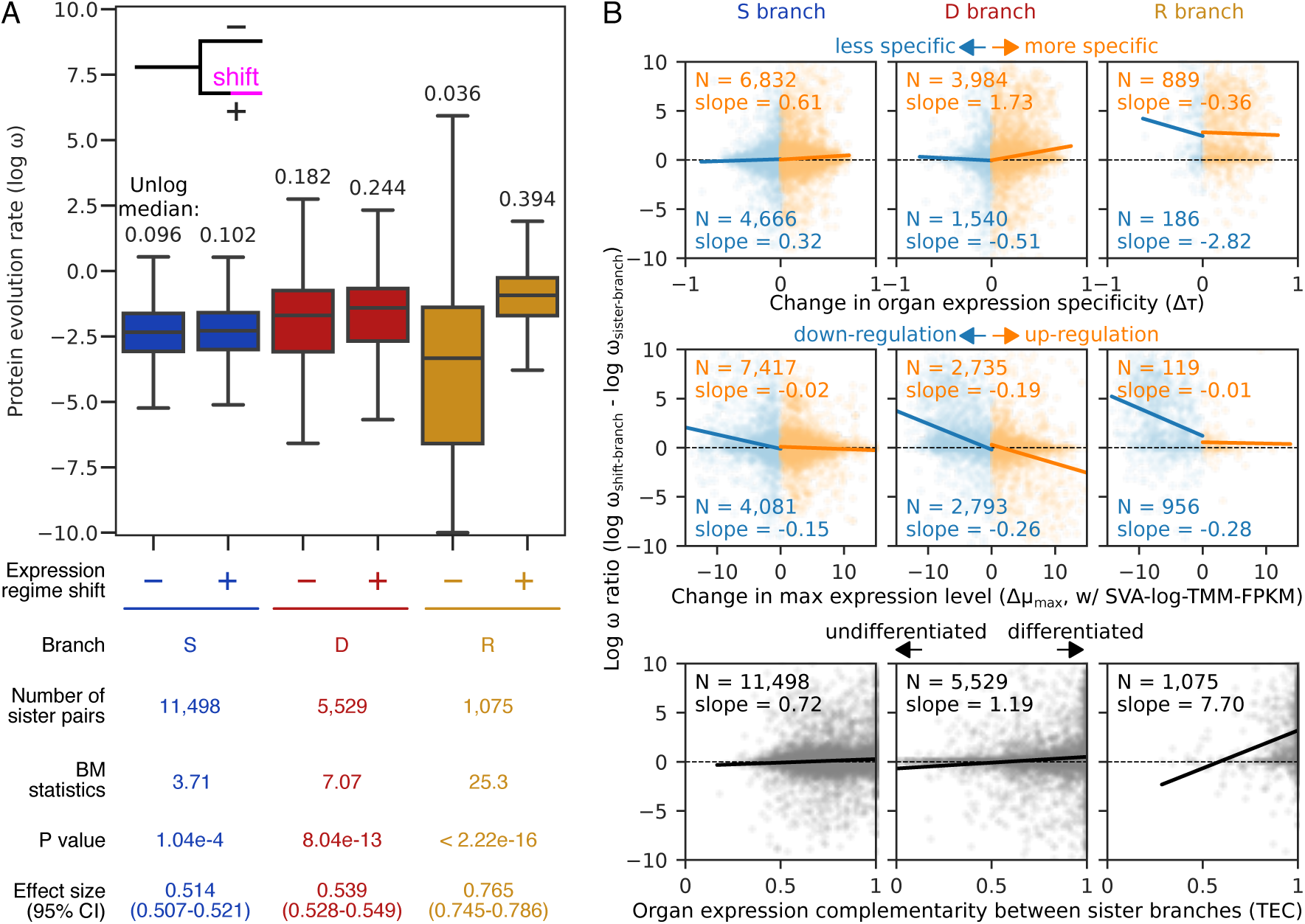
Context-dependent changes of protein evolution rate coupled with expression regime shifts. (**A**) Distribution of ω values. A plus (+) indicates branches with expression shifts, whereas minus (−) branches are sisters to the ‘plus’ branches. Statistical differences between pairs of distributions were tested using a two-sided Brunner–Munzel test (Brunner and Munzel, 2000). Non-log-transformed median values are shown above the boxplots. For visualization purpose, extreme values exceeding ± 10 were clipped. (**B**) Relationships between protein evolution rate and change in expression properties. Points correspond to expression regime shifts. Dashed lines indicate no between-branch difference in ω. Solid lines show a linear regression. Its slope and number of regime shifts are provided in the plot. Regime shifts with negative and positive changes were separately analyzed for organ expression specificity (upper) and maximum expression level (middle).

Although 52.3% of all branches with expression regime shifts had higher ω relative to sister branches (ω ratio), 27.5% of sister branch pairs are relatively undifferentiated, with differences in ω within ±5%, and 42.5% had lower ω in the branches with expression shifts. This finding led us to hypothesize that the direction of the rate changes in protein evolution is linked to how, rather than whether, expression is changed. Strikingly, we found a context-dependent association between protein and expression evolution (Fig. 4B; Fig. S9B). Increased ω ratio was linked strongly to increased, rather than decreased, organ expression specificity in S and D branches, potentially reflecting adaptive evolution coupled with specialized expression. However, it was in turn highly associated with decreased specificity in R branches, which may be explained by frequent gene decay in unsuccessful retro-copies. The change in maximum expression level was overall negatively correlated with ω ratio, but this link was stronger in down-regulation compared with up-regulation, except for D branches where the ω ratio was smaller when Δμ_max_ was larger (Fig. 4B). It has been reported that high expression slows protein evolution (Zhang and Li, 2004), and our results suggest that DNA-based duplication creates such constraints when accompanied by up-regulation. The organ expression complementarity between sister lineages was positively correlated with ω ratio (Fig. 4B), and its association was strongest in R branches, suggesting that protein evolution accelerates as gene expression patterns differentiate from their ancestral state. Collectively, these results suggest that protein evolution rate is linked to changes in expression properties through a complicated association, which masks their relationships in a global, unstratified analysis, and potentially explain a previous report of no strong relation (Warnefors and Kaessmann, 2013).

### Organ-specific propensity in gene expression evolution

The preadaptation hypothesis predicts that the ancestral organ expression prior to the shifts will affect which organs are likely to become the target of newly specific expression. To assess this, we tested whether expression shifts are random with respect to change in expression from one organ to another, by characterizing the organ in which genes are most highly expressed (PEO, primary-expressed organ).

Across vertebrates, switching from one PEO to another was detected in 6,886, 3,586, and 746 regime shifts in S, D, and R branches, respectively. The gain/loss ratios are heterogeneous among organs (Fig. 5A), suggesting that vertebrate organs serve as both sources and sinks in expression evolution, but that their relative contributions are organ-specific. Although S, D, and R branches shared a global trend of relatively abundant testis-related PEO shifts, their distributions are largely different. D branches were moderately similar to both S and R branches (Spearman’s ρ ∼ 0.6), but S and R branches were dissimilar (ρ = 0.28). This pattern, including the abundant shifts related to testis, was robust against the correction by the organ-wise numbers of expressed genes (Fig. S10). This result suggests a role for gene duplication, including by retrotranspositions, in remodeling the among-organ flow of expressed genes.

**Figure 5.**
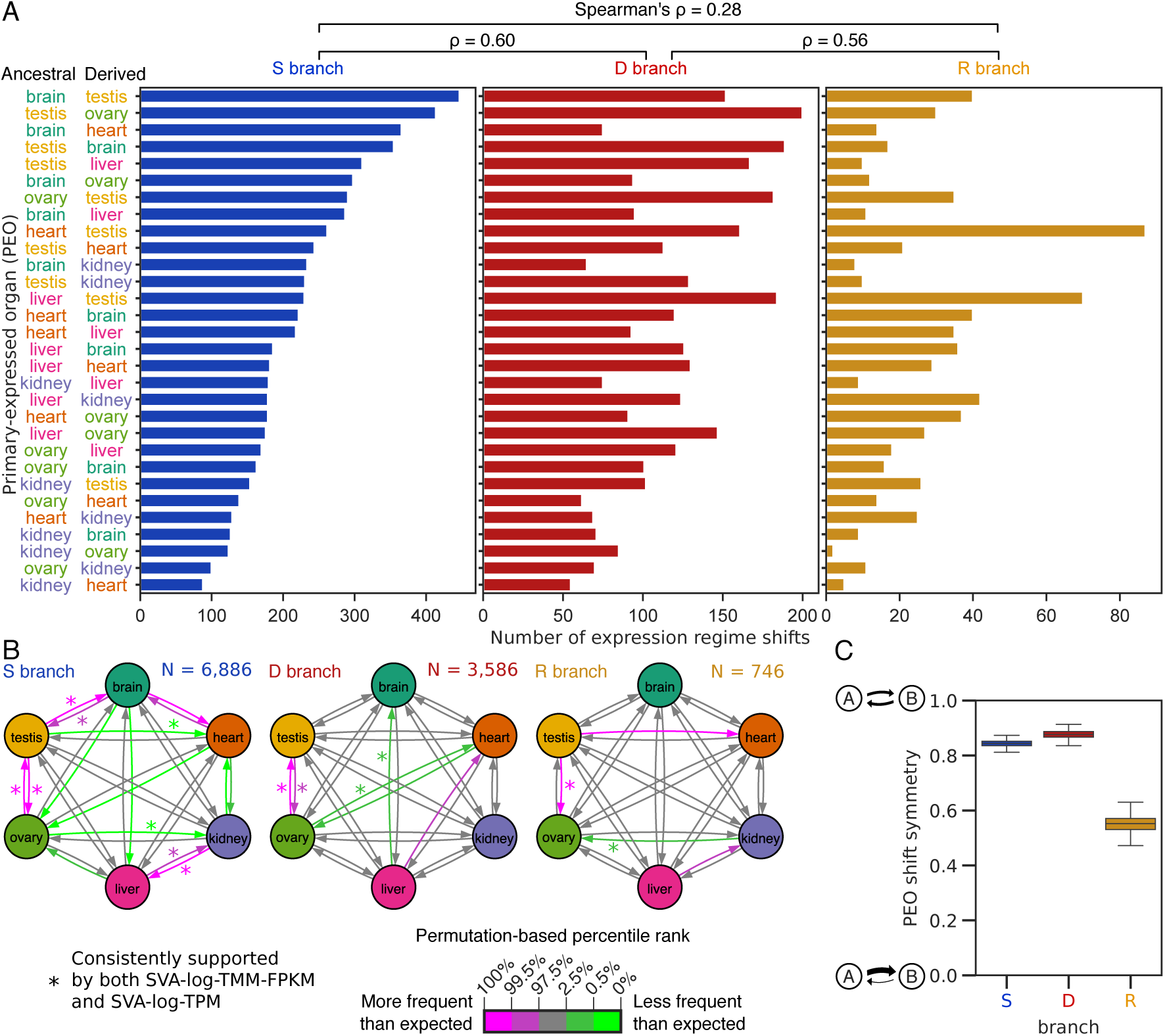
Evolutionary dynamics of gene expression. (**A**) Shift distributions of primary-expressed organs (PEOs). Y axis was sorted by abundance in S branches. Spearman’s correlation coefficients among S, D, and R branches are shown above the plots. (**B**) Preadaptation networks in organ expression. Arrows represent transitions from ancestral PEOs to derived PEOs, and its color shows statistical significance based on 10,000 permutations. The results were obtained with SVA-log-TMM-FPKM, and an asterisk (*) indicates the statistical significance supported also by SVA-log-TMM-based analysis (Fig. S11). (**C**) The global polarity of PEO shifts. The global polarity is defined by the scaled sum of differences between two opposite PEO shifts. Boxplots show the distribution estimated by 1,000 bootstrap resampling.

Controlling the total number of shifts from and to each PEO, some PEO shifts are significantly different from the random expectation (Fig. 5B). There are clear patterns of evolutionary transitions that are statistically supported by independent OU modeling of the two expression metrics. In S branches, the pairs of brain–testis and testis–ovary showed strong connections, indicating a solid exchange module. Kidney and liver also donate genes to one another, forming a separate module from brain–testis–ovary. D and R branches showed a pronounced acceleration of PEO shifts between testis and ovary. PEO shifts in S and D branches were moderately symmetric in the flow between pairs of organs (Fig. 5C), meaning two organs tend to donate comparable numbers of expressed genes each other. In contrast, R branches were more asymmetric. The lower symmetry may have perturbed the evolutionary dynamics of gene expression.

To check the robustness of our analysis, we analyzed high-confidence subsets of expression regime shifts. The most drastic expression changes were characterized by introducing a cutoff of organ expression specificity (τ > 0.5) to define organ-specific genes. Although some previously significant trends were not recovered due to small sample sizes, the result was largely consistent with the broader analysis (Fig. S11). Because tree inference errors can bias the downstream analysis including OU modeling, we also analyzed expression regime shifts found in clades which have a high support in tree inference (>99% bootstrap support). Again, the results were largely consistent (Fig. S11). Especially, the brain–testis–ovary and kidney–liver modules in S branches and the testis–ovary connection in D and R branches were always reproduced in the analyses with above thresholds in combinations with the two expression metrics (SVA-log-TMM-FPKM and SVA-log-TPM; Fig. S11), demonstrating the robustness of our conclusion.

The effect of gene trees was further examined by replicating the analysis with alternative tree topologies inferred by species tree reconciliation, which takes into account duplication–loss rates (Morel et al., 2019). This reconciliation step is expected to correct erroneous tree topology, while possibly introducing another bias derived from over-correction of biological signals such as incomplete lineage sorting. With the reconciled trees, the OU modeling with SVA-log-TMM-FPKM values resulted in equivalent numbers of expression shifts in S and D branches compared with those with non-reconciled trees (97% [23,231/23,985] and 104% [9,407/9,018], respectively) (Fig. S12A). In contrast, the phylogeny reconciliation substantially reduced the number of shifts in R branches (39% [481/1,238]). This could be explained by the correction of erroneous tree topology caused by the fast-evolving retro-copies (Fig. 4A), although the differences in shift numbers did not correlate with the topological differences measured by Robinson-Foulds distance (Robinson and Foulds, 1981) (Fig. S12B). Nevertheless, resulting PEO shift distributions were largely similar (Fig. S12C), with the reproduced accelerations in the brain–testis–ovary and kidney–liver modules (Fig. S12D) and the asymmetric PEO shifts in R branches (Fig. S12E), suggesting the robustness of detected modules against gene tree topology.

To obtain insight into the biological relevance of the among-organ modules, we characterized human genes involved in PEO shifts from all branch categories using the Kyoto Encyclopedia of Genes and Genomes (KEGG) pathway enrichment analysis. The genes descended from the testis–ovary PEO shifts enriched only one KEGG pathway term “Cell cycle” (Table S5; adjusted *P* value < 0.05), likely reflecting their function in meiosis. Although the adjusted *P* value was not statistically significant, it is noteworthy that the top-ranked term for the brain–testis connection was “Endocrine and other factor-regulated calcium reabsorption” annotated to four genes including GNAQ, which has been implicated to tumor formation in neuronal tissues (Gessi et al., 2013; Van Raamsdonk et al., 2009). In the kidney–liver module, 15 terms were significantly enriched, many of which appear to be related to the functions and diseases in those organs, for example, “Bile secretion”, “Phagosome”, “Lysosome”, “ABC transporters”, “Sphingolipid metabolism”, and “Type I diabetes mellitus” (Table S5). These results suggest that the among-organ modules in the PEO shifts played a role in supplying functionally important genes.

Because ancestral expression was shown to orient new expression by the analysis of PEO shifts, we concluded that there was prevalent organ-specific propensity, which supports the presence of preadaptation in gene expression.

## Discussion

Our results suggest that the landscape of expression evolution is strongly shaped by mechanisms of gene birth. Expression shifts are more pronounced following gene duplication in agreement with the results of pairwise gene expression analyses (Assis and Bachtrog, 2015; Guschanski et al., 2017; Huerta-Cepas et al., 2011), and shifts in patterns of primary-expressed organs strongly depend on the expression state in the ancestral organism. Thus, by analyzing such influences on a genome-wide scale for a moderately large number of species, the question whether long-term expression in one organ predisposes genes to be subsequently utilized in other organs has been answered in the affirmative. There are preadaptive propensities in the evolution of vertebrate gene expression, and the propensity varies with the presence and type of gene duplication. Furthermore, the approach developed in this study, using complex gene family phylogenies including gene duplications and losses that do not assume perfect match to the species phylogeny, and incorporating a curation pipeline to amalgamate large amounts of transcriptome data from many studies, was essential to obtain the necessary species density and phylogenetic resolution to answer this question. The extensibility of this method will allow for more species and more organs to be incorporated as further studies come into the literature from diverse laboratories.

The mechanisms responsible for the preadaptive propensities that influence expression shifts among organs are, however, unknown. A key question in understanding these shifts may be the role of adaptation in the shift, and in subsequent evolution. We have been careful so far to simply describe the shifts, but adaptive possibilities include subfunctionalization, escape from adaptive conflict (EAC), and neofunctionalization (Conant and Wolfe, 2008; Des Marais and Rausher, 2008; Sikosek et al., 2012). The increased number of shifts following duplication suggests that drift alone is not the explanation, but subfunctionalization easily could be. Subfunctionalization is the idea that, if a gene has multiple functions prior to duplication, they may be segregated among the duplicates following gene duplication. Thus, the expression shifts may be simply a shift in focus of a duplicated copy on a subset of the necessary expression profile needed at the organismal level. In this scenario, any accompanying acceleration of amino acid substitution would be caused by a loss of constraint and reduced purifying selection in one expression environment or the other.

EAC involves more adaptation by adding the simple idea that prior to duplication, the multiple functions and expression regimes were at least partially in conflict. Such conflicts could clearly occur at the amino acid level, but could also occur at the expression level. For example, if expression levels were focused on a most-important tissue or most-sensitive tissue prior to duplication, but after duplication could be more tailored to what is better for the new expression regime. Finally, neofunctionalization would occur at the sequence or expression level, if the loss of selection on a duplicate allowed mutations that were previously harmful to the old function, but now are not, and are able to carry out some novel functional aspect that was previously prohibited. Neofunctionalization is perhaps the most interesting and extraordinary possible cause for the expression regime shifts we see, but it requires strong evidence and it is not a necessary explanation for what we observe.

In this context, the patterns of expression regime shifts we observed may be explained at different levels of biological organization, from the tissues and cells that make up organs, to subcellular compartments, chromatin structure, promoter usage, and protein biochemistry. Part of the propensity shifts we observed can be explained by the “out of the testis” hypothesis, which posits that accelerated gains of testis expression are based on the permissive chromatin state, abundant transcriptional machinery, relatively simple promoters required for the expression in spermatogenic cells and following gains of new expression patterns (Kaessmann, 2010; Kleene, 2005). This theory fits to the accelerated testis-related PEO shifts, and could fit with any of the adaptive scenarios discussed above, but the other detected patterns (e.g., kidney–liver module) require other explanations.

One potential mechanism of preferences in expression regime shifts is a cell-type or sub-cellular component mechanism. In such a mechanism, if two organs tend to share cell types or usage of sub-cellular components, they may be prone to expropriate genes between the two organs. It is known that gene expression levels in the kidney and liver tend to change jointly, possibly reflecting their similar physiology including detoxifications and waste excretion (Brawand et al., 2011). Such functional similarity may also explain the presence of the kidney–liver module of gene exchange.

Another possibility is a regulatory mechanism whereby frequent gene-exchanging organs use similar sets of regulatory elements. Altered expression between such organs could occur with relatively few mutations in regulatory sequences. *Cis*-regulation is indeed a major source of expression evolution, as it explains a certain fraction of expression variability, for example, in budding yeast (30% in duplicates and 19% in singletons) and undergoes a more rapid divergence than *trans*-regulation (Dong et al., 2011).

Finally, another possible mechanism for expression regime shifts following gene duplication is at the protein level. If frequently interacting organs have similar environmental requirements for expressed proteins, a few amino acid substitutions may tend to be required to adjust biochemical properties. Protein reusability may be determined by cellular environments such as pH and temperature or by functional categories of proteins. Our analysis indicates that the regime shifts that drastically differentiate the expression tend to be coupled with accelerated protein evolution (Fig. 4B), and this result can be viewed as a support for a protein-level mechanism. In such a mechanism, synergistic resolution of EAC may be a driving force for changes in both amino acid composition and expression regimes. We note that these mechanistic hypotheses are not mutually exclusive, and varying combinations of factors may contribute to generate preadaptive patterns of gene expression.

In this study, we established a method to standardize RNA-seq data from disparate research projects and developed a pathway for data amalgamation. Thanks to multiple rounds of innovations in sequencing technology, transcriptome data are being produced at an unprecedented rate in a greater variety of organisms and samples, such as those for multi-species multi-organ developmental series (Cardoso-Moreira et al., 2019). The transcriptome amalgamation will expand the use of such resources to study gene expression evolution.

By reconstructing gene expression in gene family phylogenies, our analysis revealed non-randomness and directionality of expression evolution. This suggests prevalent preadaptation in gene expression, and that adaptation to expression in certain organs is more conducive to future expression in other organs. This provides further details on how gene duplication has helped to reshape the dynamics of expression evolution that contributed to the vertebrate diversification.

## Materials and Methods

### Species selection

A total of 105 species included in the Ensembl release 91 (Yates et al., 2016) were searched for data availability in the NCBI SRA database (Leinonen et al., 2011) (final search on May 1, 2018) and 22 species were found to have RNA-seq data for 6 organs: brain, heart, kidney, liver, ovary, and testis. *Lepisosteus oculatus* was excluded due to an insufficient quality of available expression data, and therefore remaining 21 species were selected for further analysis.

### Species tree

The dated species tree for the 21 species was retrieved from TimeTree (Hedges et al., 2006) (downloaded on March 15, 2018; Supplementary Data). Some species were unavailable in the database and therefore they were temporarily replaced by closely related species to obtain the dated species tree.

### Gene sets

Coding sequences (CDS) were retrieved from the Ensembl database. The longest transcript was retained when multiple transcripts were annotated to the gene. The quality of gene sets was evaluated using BUSCO 4.0.5 (Simão et al., 2015) with the single-copy ortholog set “vertebrata_odb10” (Fig. S8).

### Transcriptome metadata curation

We developed an automated python program for SRA metadata curation (Table S3; Supplementary Data). RNA-seq data were selected from the NCBI SRA database by keyword searches limited to the 21 species, the 6 organs, and Illumina sequencing platforms. Orthographical variants of annotations were standardized with keyword libraries created by manually checking the original annotations. Prenatal or unhealthy samples and small-scale sequencing samples (<5 million reads) were excluded. Data for non-messenger RNA sequencings were also removed. In treatment and control RNA-seq pairs, only control experiments were included.

### Transcriptome quantification

Fastq files were extracted from downloaded SRA files using parallel-fastq-dump 0.6.2 (https://github.com/rvalieris/parallel-fastq-dump) with the minimum read length of 25 bp and the quality filter (-E option) (Leinonen et al., 2011). The fastq sequences were then subjected to a quality filtering by fastp 0.12.3 (Chen et al., 2018). The filtered reads were mapped to genomic features annotated as non-messenger RNAs in the Ensembl GTF files using bowtie2 2.3.4 (Langmead and Salzberg, 2012) and resultant unmapped reads were used for expression level quantification using kallisto 0.43.1 with the sequence-based bias correction (Bray et al., 2016). Samples were removed if 20% or smaller percentages of reads were mappable (Fig. S2). Estimated mapped read counts and transcript lengths were used to calculate TPM and FPKM values. For the latter, the “trimmed mean of M-values (TMM)” normalization method was applied (Robinson and Oshlack, 2010). Sample-wise TMM scaling factors were obtained across all RNA-seq samples using the FPKM values of 1,377 single-copy orthologs. Because TMM and TPM transformations are incompatible with each other, the TMM scaling was applied only to FPKM values. TPM and TMM-FPKM values were subsequently transformed to *log*(*N* + 1) values (log-TPM and log-TMM-FPKM, respectively). Paralogous genes that haven’t diverged in their nucleotide sequences could not be distinguished well in the quantification step. Although our scope is to characterize gene expression in the timescale of vertebrate evolution, this difficulty likely leads to an underestimation of expression regime shifts in young duplicates.

### Iterative anomalous sample removal coupled with surrogate variable analysis

Anomalous RNA-seq samples were iteratively removed by a correlation analysis coupled with an expression bias correction. In each iteration, unwanted biases in expression level were removed by surrogate variable analysis (SVA, ‘sva’ function in an R package ‘sva’) (Leek and Storey, 2007). SVA analysis was applied at the beginning of each iteration so that it is not influenced by anomalous samples removed in previous iterations. Subsequently, Pearson’s correlation coefficients were calculated for every RNA-seq data against mean expression level in each organ generated by averaging all other data excluding those from the same BioProject (Fig. S3). We assume that the sample’s correlation coefficient against the same organ is higher than any of the values against the other organs, and we removed all samples from the same BioProject when violations were found. These steps were repeated until no violations were left and SVA-corrected expression levels were finally reported (SVA-log-TMM-FPKM and SVA-log-TPM). The curation steps were skipped if multiple samples were unavailable in the species and hence SVA analysis was inapplicable. The final dataset was comprised of 1,903 RNA-seq experiments from 182 BioProjects that cover 6 organs from 21 vertebrate species without missing data (Tables S1–S2).

### Orthogroup classification

Orthogroups, which contain all genes descended from one gene in the common ancestor, were inferred from coding sequences of the 21 species using OrthoFinder 2.1.2 (Emms and Kelly, 2015) guided by the species tree. In total, 17,896 orthogroups were generated. The largest orthogroup, which is comprised of 7,893 olfactory receptor genes, was removed from the analysis because of computational burden. After sequence alignment processing (see “Multiple sequence alignment”), we removed small orthogroups which retained less than four genes and orthogroups that showed no parsimony informative sites, because phylogenetic relationships cannot be inferred. As the result of filtering, 15,280 orthogroups were left for OU modeling, with the largest one containing 3,796 zinc-finger proteins (Table S4).

### Multiple sequence alignment

Multiple fasta files containing coding sequences were generated for each orthogroup. Stops and ambiguous codons were masked as gaps (for implementation, see https://github.com/kfuku52/cdskit). In-frame multiple codon sequence alignments were generated using MAFFT 7.394 with the ‘auto’ option (Katoh and Standley, 2013) and tranalign in EMBOSS 6.5.7.0 (Rice et al., 2000). Anomalous genes were excluded by MaxAlign (Gouveia-Oliveira et al., 2007) which decreased the largest orthogroup size from 3,796 to 2,382 genes. Spurious codons were removed in-frame using pgtrimal in Phylogears2-2.0.2016.09.06 (https://www.fifthdimension.jp/products/phylogears/) with the ‘gappyout’ option (Capella-Gutierrez et al., 2009).

### Gene tree reconstruction

Maximum-likelihood trees were reconstructed using IQ-TREE 1.6.5 (Nguyen et al., 2015) with the best-fit nucleotide substitution models selected by ModelFinder with the Bayesian Information Criterion (Kalyaanamoorthy et al., 2017). Larger orthogroups and longer genes tended to fit more complex substitution matrices and larger numbers of categories of rate heterogeneity (Fig. S13A–B; Table S4). Ultrafast bootstrapping with 1,000 replicates was performed to evaluate the credibility of tree topology (Minh et al., 2013) with a further optimization of each bootstrapping tree (-bnni option) (Hoang et al., 2018). To evaluate the effect of alternative gene tree topologies, we performed phylogeny reconciliation using GeneRax 1.0.0 (Morel et al., 2019) with the maximum-likelihood gene trees and the species tree as input. Because rooted trees were generated in this step, the tree rooting (described below) was skipped for the reconciled trees.

### Reconciliation-assisted gene tree rooting

Candidate rooting positions were inferred with different methods. Using the dated species tree, all rooting branches with the minimum duplication-loss score were identified using the rooting mode of NOTUNG 2.9 with the default parameters (duplication score = 1.5, loss score = 1.0) (Chen et al., 2000). The midpoint of the longest path (Farris, 1972) and the position with the minimal ancestor deviations (MAD) (Tria et al., 2017) were also considered as candidates. The final rooting position was reported based on overlaps among those rooting positions (Figs. S13C,E; Table S4).

### Reconciliation-assisted divergence time estimation

To prepare dated gene trees, we first matched species tree nodes with corresponding gene tree nodes using the reconciliation mode of NOTUNG 2.9 (Chen et al., 2000) and created time constraints of speciation nodes (Fig. S13F). Duplication nodes were constrained with the upper and lower age limits derived from corresponding speciation nodes. If the root node is a duplication node and is not covered by the range of the species tree, the upper age limit was set to 1,105 million years ago, which corresponds to the split of animals and fungi (Hedges et al., 2006). Divergence time was then estimated by a penalized likelihood method (Sanderson, 2002) implemented in an R package ‘ape’ (‘chronos’ function with ‘discrete’ model) (Popescu et al., 2012) with time constraints on speciation, duplication, and root nodes. When reasonable initial parameters were not found after 1,000 trials, the above constraints were partly relaxed (Fig. S13D; Table S4). The implementation is provided on GitHub (https://github.com/kfuku52/RADTE).

### Modeling and shift detection of expression evolution

Using the dated gene trees and organ-wise mean values of SVA-log-TMM-FPKM and SVA-log-TPM, regime shifts in gene expression were detected as shifts of optimal trait values in Ornstein–Uhlenbeck (OU) models determined by a Lasso-based model selection with AICc in an R package ‘*l*1ou’ (Khabbazian et al., 2016). Because there is no available software to handle within-species variation in phylogenetic OU shift detection without predefined hypotheses on the number and place of regime shifts, we used mean expression level as the input. It is shown by simulation that the species mean and species variance models show comparable power in the regime shift detection (Rohlfs et al., 2014), suggesting that our species mean model is expected to perform as good as the species variance model. In the model, α and σ^2^ parameters were assumed unchanged in the tree (Khabbazian et al., 2016), and therefore only the global, rather than branch- or clade-wise, stationary variance (γ) were obtained. Expression levels in the six organs were treated as multivariate traits where α and σ^2^ were estimated for each organ but regime shifts were assumed to occur jointly in the same set of branches (Khabbazian et al., 2016). The phylogenetic mean (expression level at the root node) was estimated with the “OUfixedRoot” model. To handle gene trees recalcitrant to this analysis (especially those with a large number of genes), we skimmed gene trees by collapsing clades with small changes in expression level (Fig. S14). Specifically, we first calculated all-vs-all Pearson’s correlation coefficients of gene expression level among all genes that belong the clade. The clade was collapsed into a single tip if the expression patterns were almost identical (minimum correlation coefficient between genes > 0.99). Phylogenetic means of the collapsed clade were calculated by assuming the Brownian motion and were used as expression level at the new tip. The upper limit of regime shifts was set as 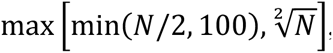, except for the largest tree with 2,382 genes where the upper limit was decreased to 10 to cope with an unrealistically large number of branch combinations to consider. The number of detected regime shifts was always smaller than the upper limit in the SVA-log-TMM-FPKM analysis, whereas three out of 15,820 trees reached the upper limit in the SVA-log-TPM analysis (Fig. S6A).

### Analysis of expression patterns

We characterized expression patterns of extant and ancestral genes by calculating different metrics from fitted values (μ) in the OU models. Organ specificity was measured by τ (Yanai et al., 2005), which outperformed other methods in a benchmark for tissue specificity (Kryuchkova-Mostacci and Robinson-Rechavi, 2016). Expression complementarity between sister lineages was measured by the metrics called ‘TEC’ (Huerta-Cepas et al., 2011). Because μ was estimated from log-transformed expression levels, these expression metrics were calculated with unlog-transformed μ values. PEOs were defined as the organ in which the highest expression levels were observed among the six organs we analyzed.

### Estimation of protein evolution rate

Parameters for codon substitution matrix, shape parameter of discrete gamma distribution for rate heterogeneity (α), equilibrium transition/transversion rate ratio (κ), equilibrium nonsynonymous/synonymous substitution rate ratio (ω) were estimated using IQ-TREE 1.6.5 (Nguyen et al., 2015) by fitting GY+F3X4+G4 models to each gene tree. Equilibrium base composition (θ) was estimated from empirical codon state frequencies which are calculated from the alignment by counting. To obtain θ at the root node, we calculated θ at subroot nodes by taking advantage of IQ-TREE’s empirical Bayesian method for ancestral sequence reconstruction. Considering subroot branch length, a weighted average of the subroot θ values were calculated as the θ value at the root node. Using all those parameters, branch-wise nonsynonymous/synonymous substitution rate ratios were estimated by stochastic substitution mapping (*mapdNdS*) (Guéguen and Duret, 2018) using bio++ library (Guéguen et al., 2013). To examine the robustness of the *mapdNdS*-based ω estimation, we compared the results with those obtained by maximum-likelihood ω estimation by fitting MG94W9 models in HyPhy 2.3.11 (Pond et al., 2005). The two methods yielded consistent results on the effect of branching events and expression shifts (Fig. 4A; Fig. S9A), suggesting methodological robustness. We reported *mapdNdS*-based results in the main text.

### Analysis of gene structure and location

The number of introns and chromosomal location were obtained for each gene from the Ensembl gene models (GFF3 files). The intron numbers were subsequently converted to binary values that represent intronless and intron-containing states. Chromosomal locations were categorized into autosome, X chromosome, and Y chromosome. Genes from non-therian species were treated as missing data because mechanisms of their sex determination are not homologous to the mammalian XY system (Veyrunes et al., 2008). Genes from *Chinchilla lanigera* were also treated as missing data because their sequenced genomes are not anchored to chromosomes in the Ensembl release 91. The posterior probabilities of ancestral character states were inferred by the stochastic character mapping of discrete traits (Bollback, 2006) implemented in an R package ‘phytools’ (Revell, 2012). Because functional retrotranspositions (Jun et al., 2009; Marques et al., 2005) and inter-chromosomal duplications (Pace et al., 2009) are rare events relative to the timescale of the vertebrate evolution on the per-gene basis, we set the transition rate parameters to a sufficiently small value (1 × 10^-3^ per gene per million years). Since intron gain occurs few orders of magnitude less frequently than its loss caused by retrotransposition and others (Roy and Gilbert, 2005), the rate of intron gain is set to be lower (1 × 10^-4^ per gene per million years).

### Analysis of branching events

Speciation and duplication nodes (S and D/R nodes, respectively) were classified by a species-overlap method (Huerta-Cepas et al., 2007) and were mapped to the species tree on the basis of species coverages of the gene tree clades. A transition from intron-containing to intronless states with a posterior probability greater than 0.5 was classified as a retrotransposition event (R node). The branches that correspond to the original copy of a retrotransposition event were not included in R branches. Although our classification cannot detect retrotranspositions from originally intronless genes, we expect such situations would be rare because most vertebrate genes contain at least one intron (e.g., 20,160/21,242 human genes). Inter-chromosomal translocation was detected by considering chromosomal locations with the highest posterior probability as the ancestral states. Because of the difficulty in determining rooting positions of deep phylogenies, gene tree nodes older than the root node of the species tree were removed from the analysis.

### KEGG pathway enrichment analysis

Human genes that descend from the shift branches were pooled for each specific PEO shift. The gene lists were examined for enrichment against an Enrichr library “KEGG_2019_Human” (Kuleshov et al., 2016) using a python package GSEApy (https://github.com/zqfang/GSEApy). Statistically significant (adjusted *P* value < 0.05) KEGG pathway terms were reported in Table S5.

### Data visualization

Phylogenetic trees were visualized using a python package ‘ETE 3’ (Huerta-Cepas et al., 2016) and an R package ‘ggtree’ (Yu et al., 2017). A part of animal silhouettes in Fig. 3A were obtained from PhyloPic (http://phylopic.org). Box plot elements of all figures are defined as follows: center line, median; box limits, upper and lower quartiles; whiskers, 1.5x interquartile range; points, outliers. Box-plot outliers are suppressed in Figs. 3, 4, S9, S11, and S12.

### Code and data availability

Scripts, parameter values, gene expression values including SVA-log-TMM-FPKM and SVA-log-TPM, and other data used in this study are available as Supplementary Data (Mendeley Data ID available on publication).

## Acknowledgements

We acknowledge the following sources for funding: MEXT/JSPS KAKENHI 18J00178 (K.F.), Sofja Kovalevskaja programme by the Alexander von Humboldt Foundation (K.F.), and NIH R01 GM083127 (D.D.P.). Computations were partially performed on the NIG supercomputer. The authors declare no competing financial interests.

## Author Contributions

K.F. and D.D.P. jointly designed the study and wrote the paper. K.F. designed and wrote all programs and performed data analysis.

## Supplementary Materials

**Fig. S1.**
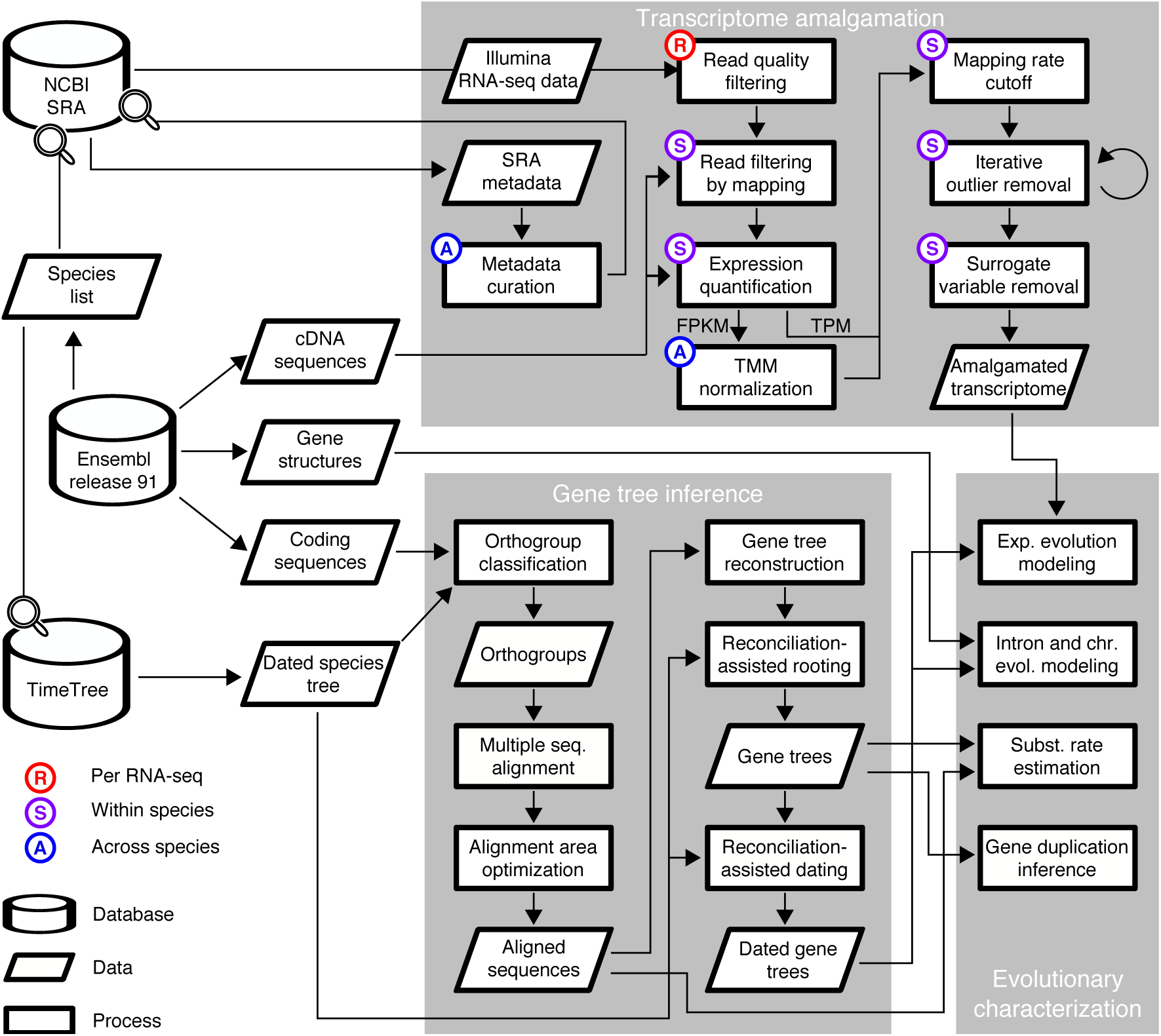
A flow-chart of transcriptome amalgamation, gene tree inference, and evolutionary characterization in this study.

**Fig. S2.**
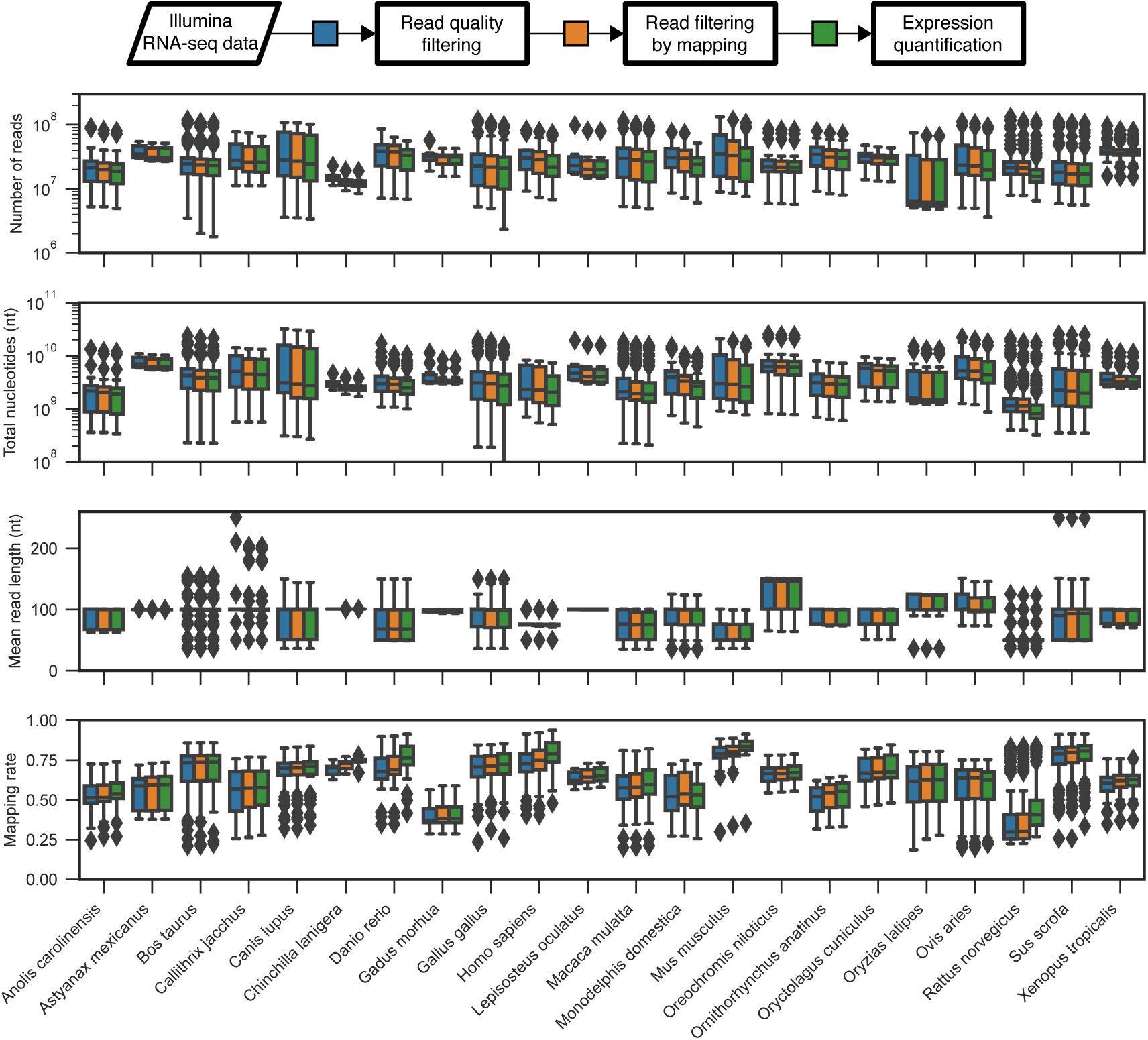
Changes in number of reads, total nucleotide length, mean read length, and mapping rate by RNA-seq read filtering. Mapping rates were, in almost every case, improved by both read quality filtering and read filtering by mapping to miscellaneous genomic features.

**Fig. S3.**
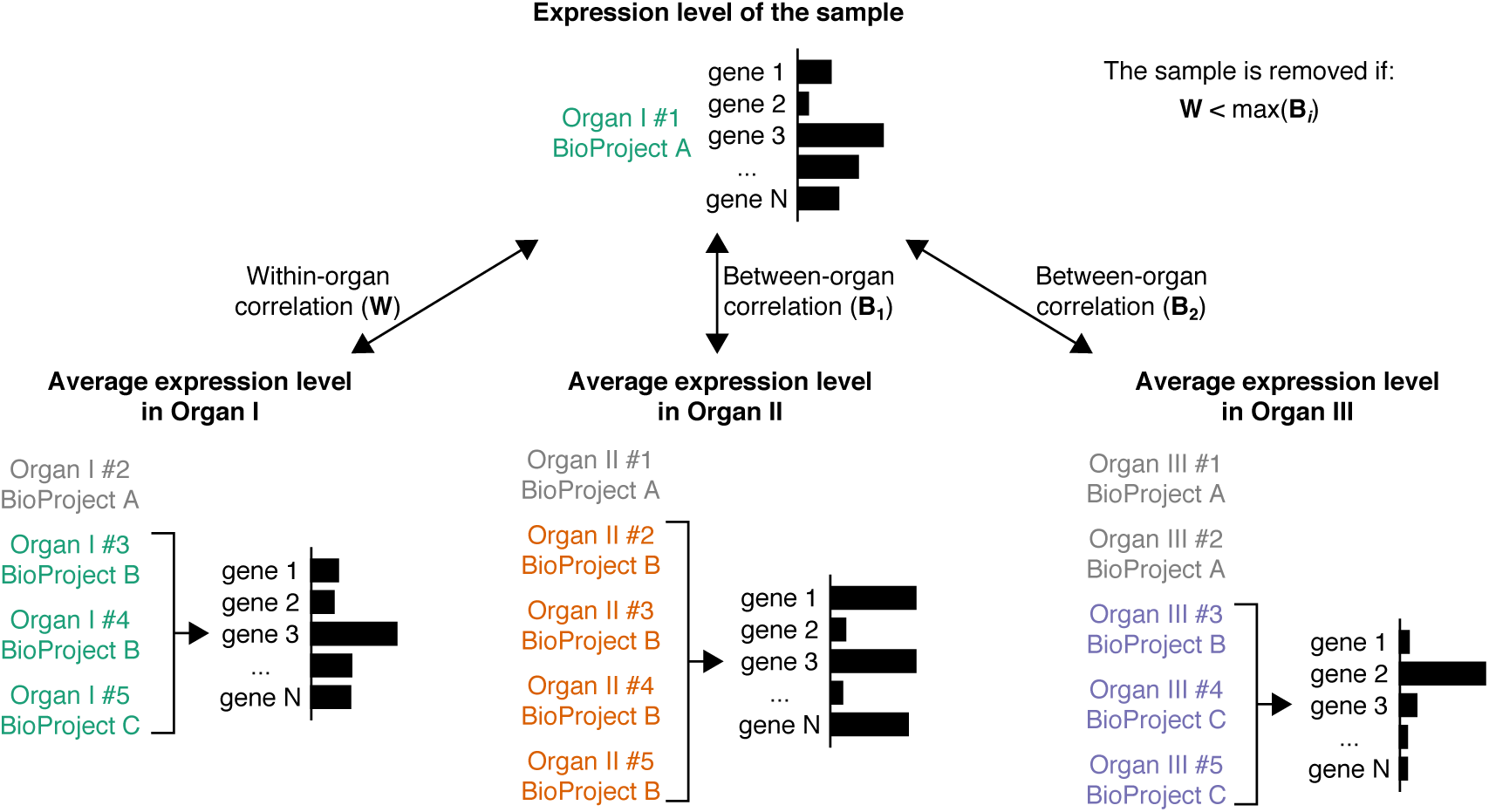
A correlation analysis for the detection and removal of anomalous RNA-seq samples. Expression levels of all genes were compared between the sample and the organ averages. A sample was removed if any between-organ comparisons yielded a correlation coefficient higher than the within-organ comparison.

**Fig. S4.**
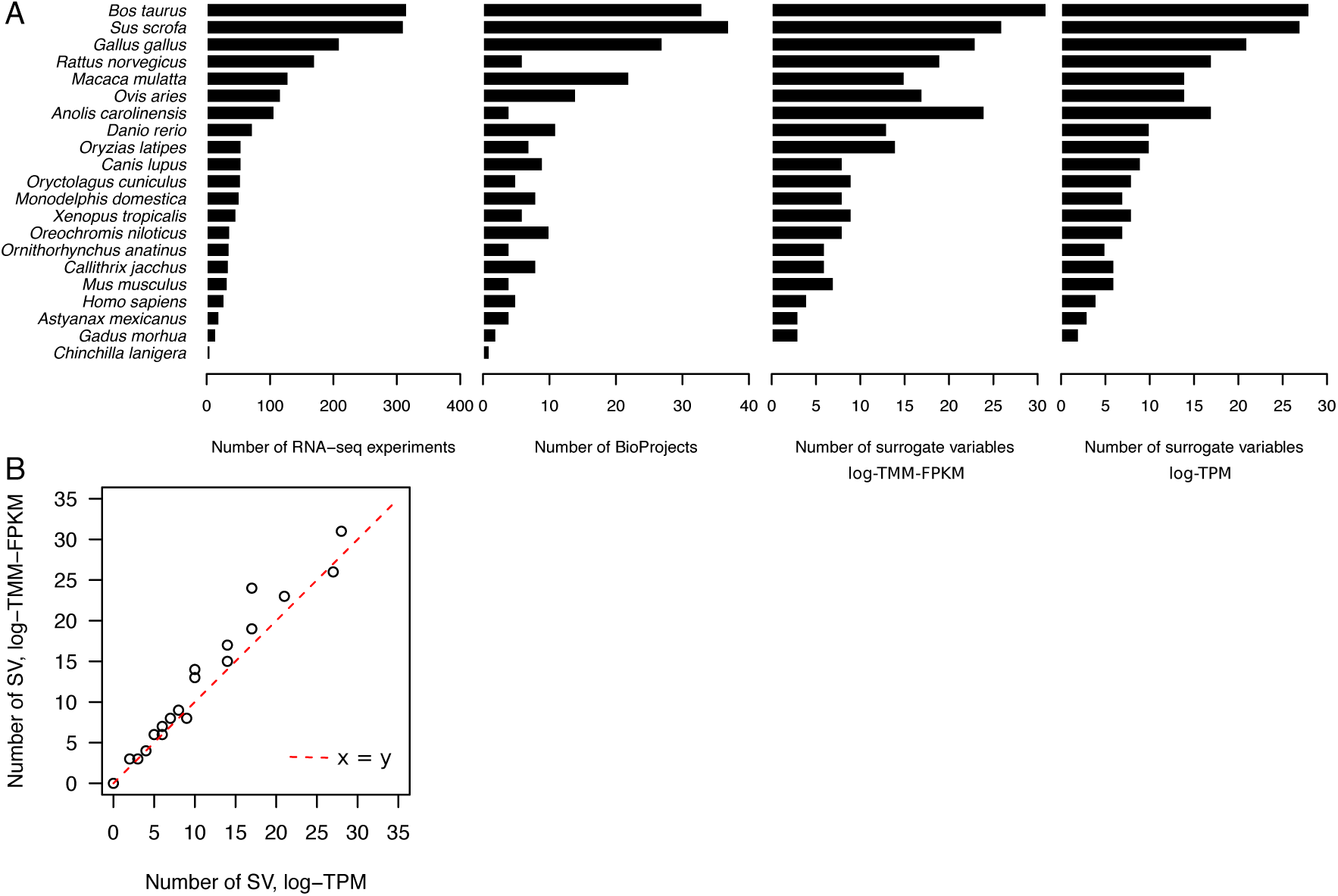
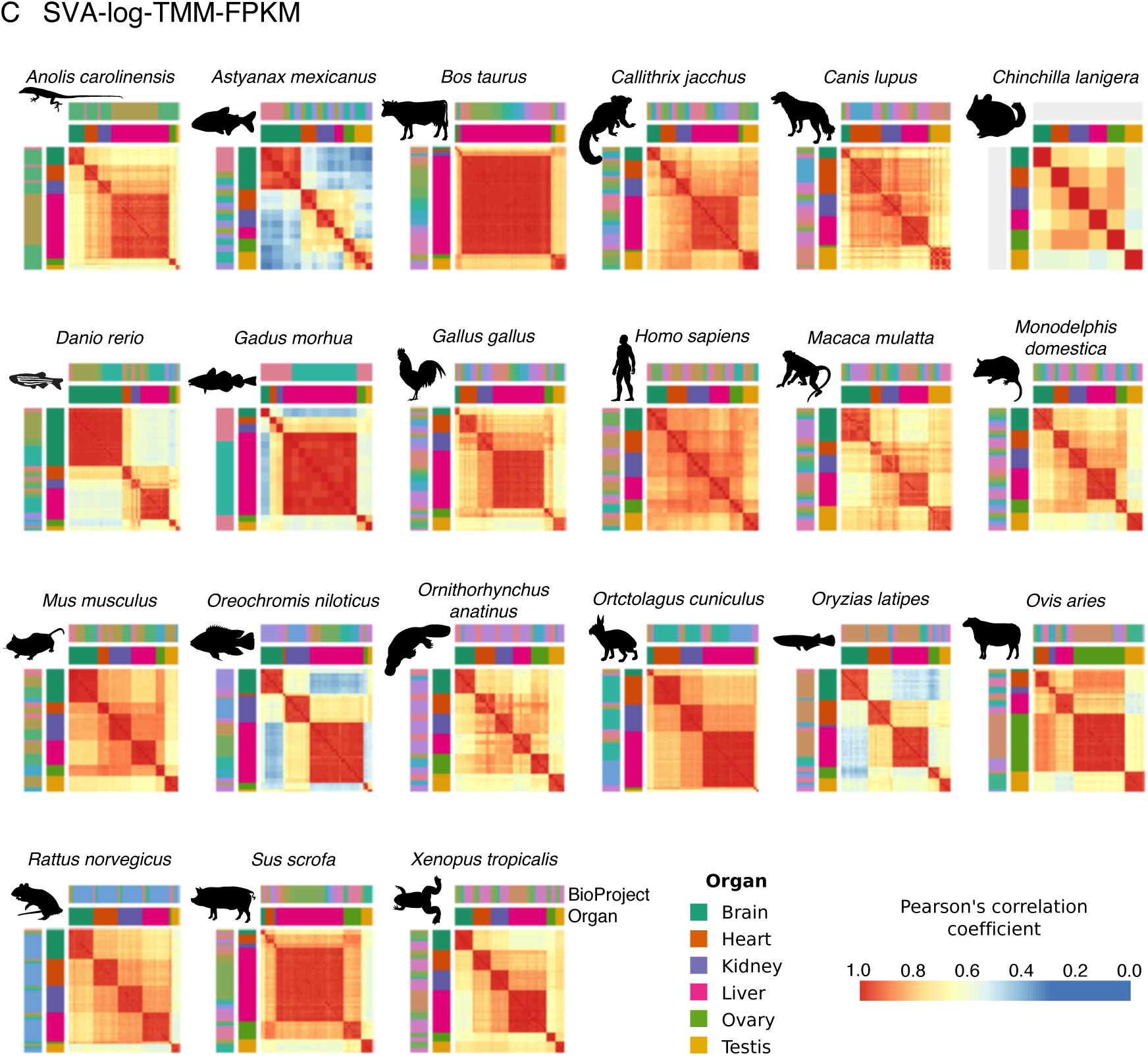
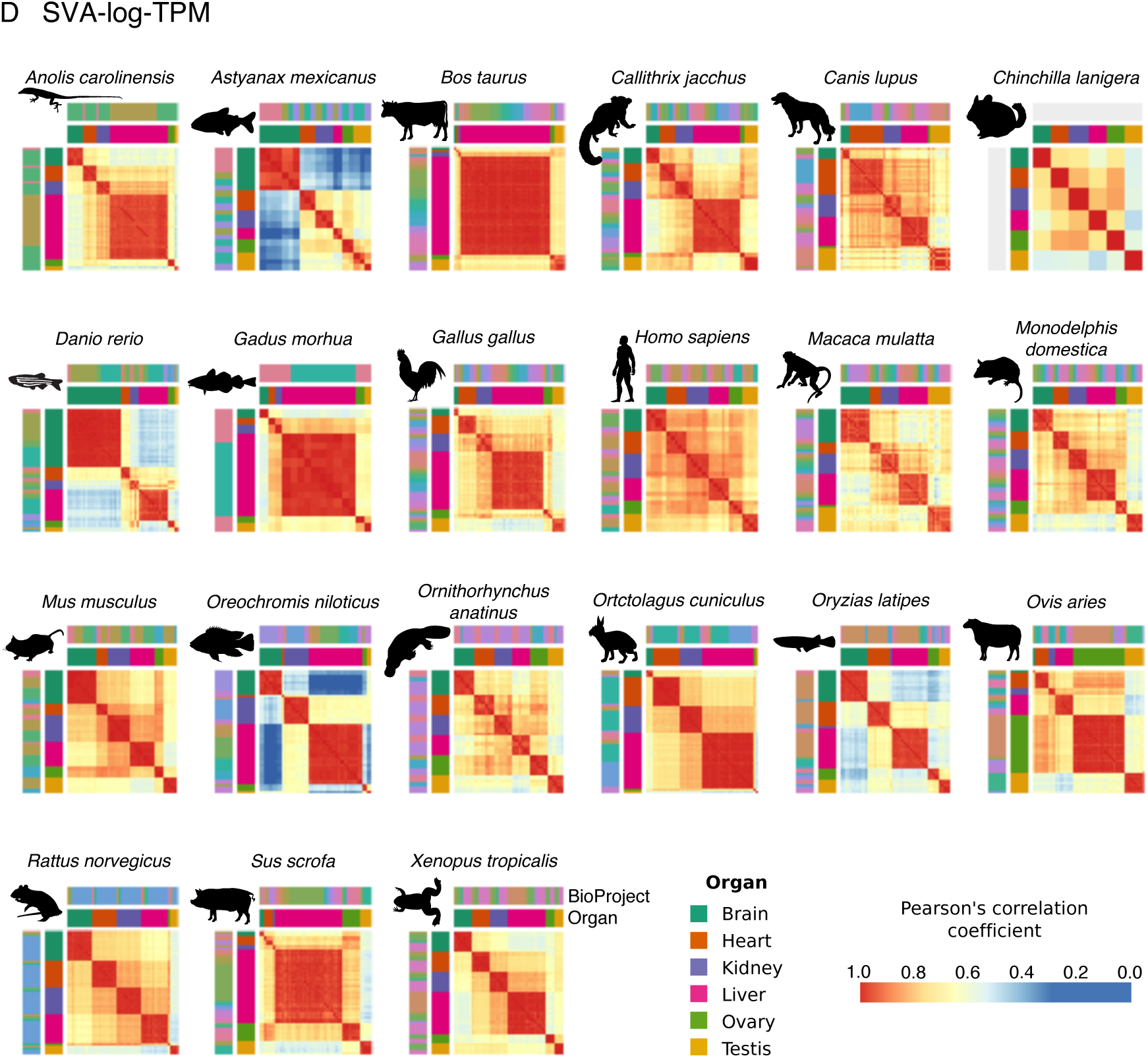
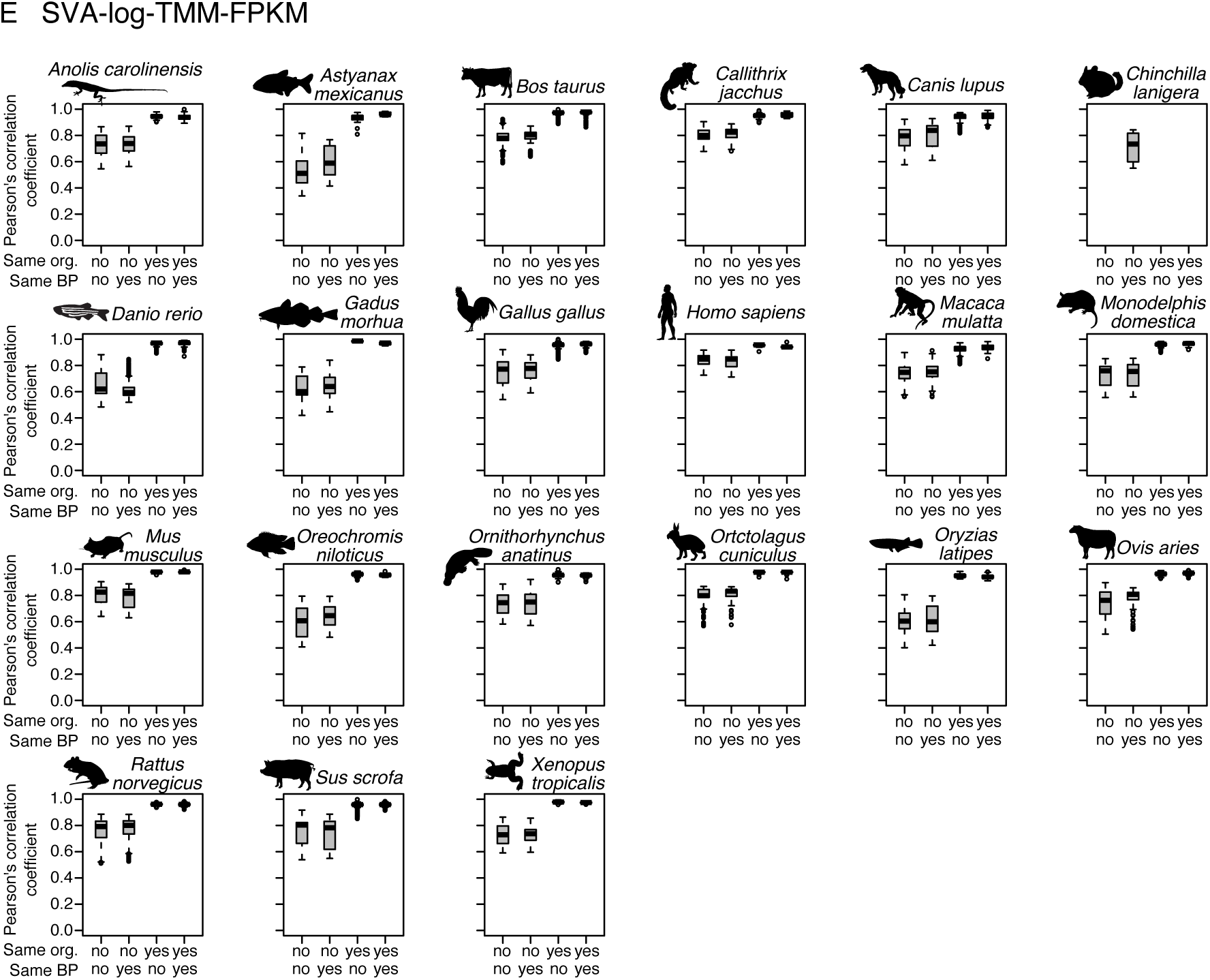
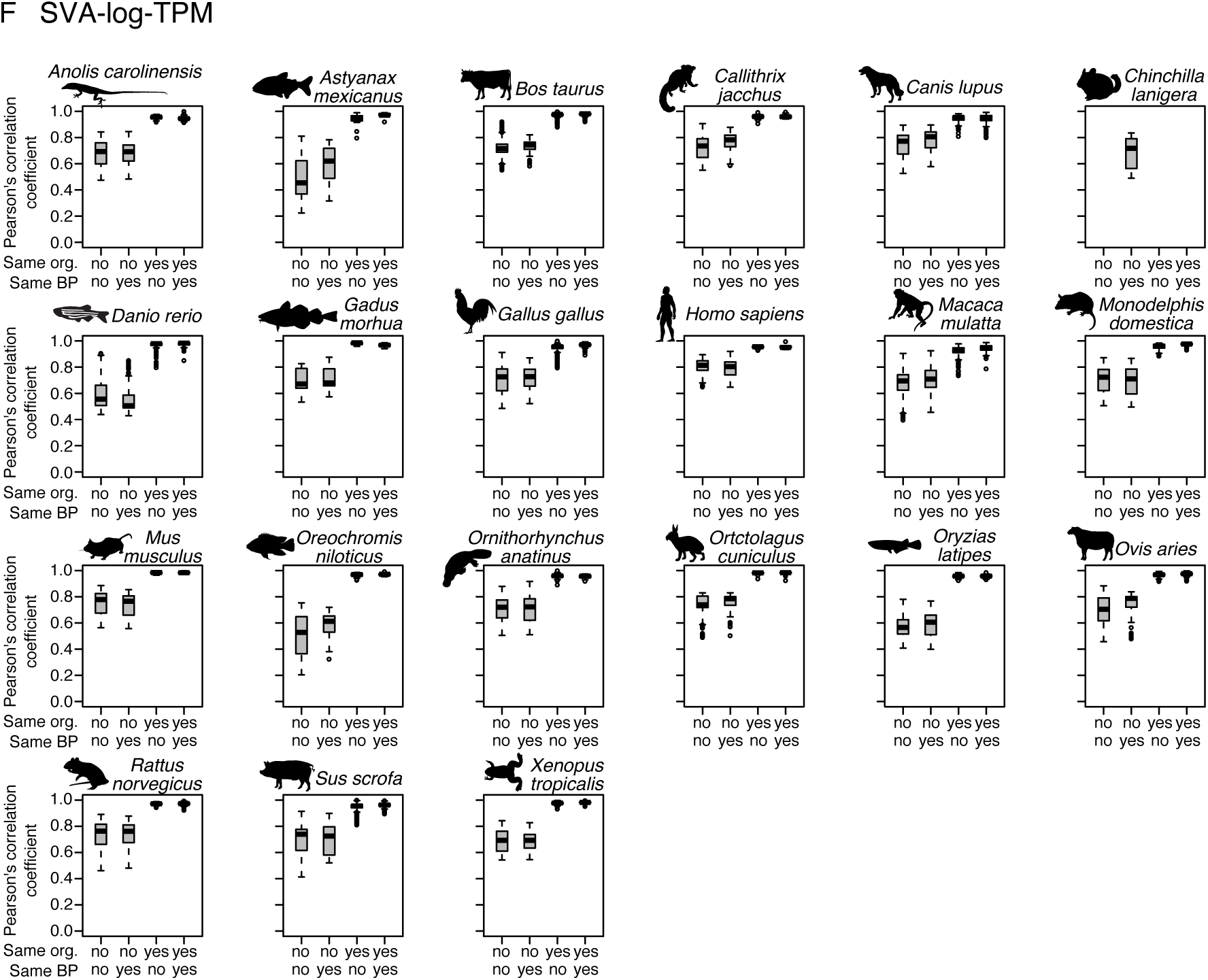
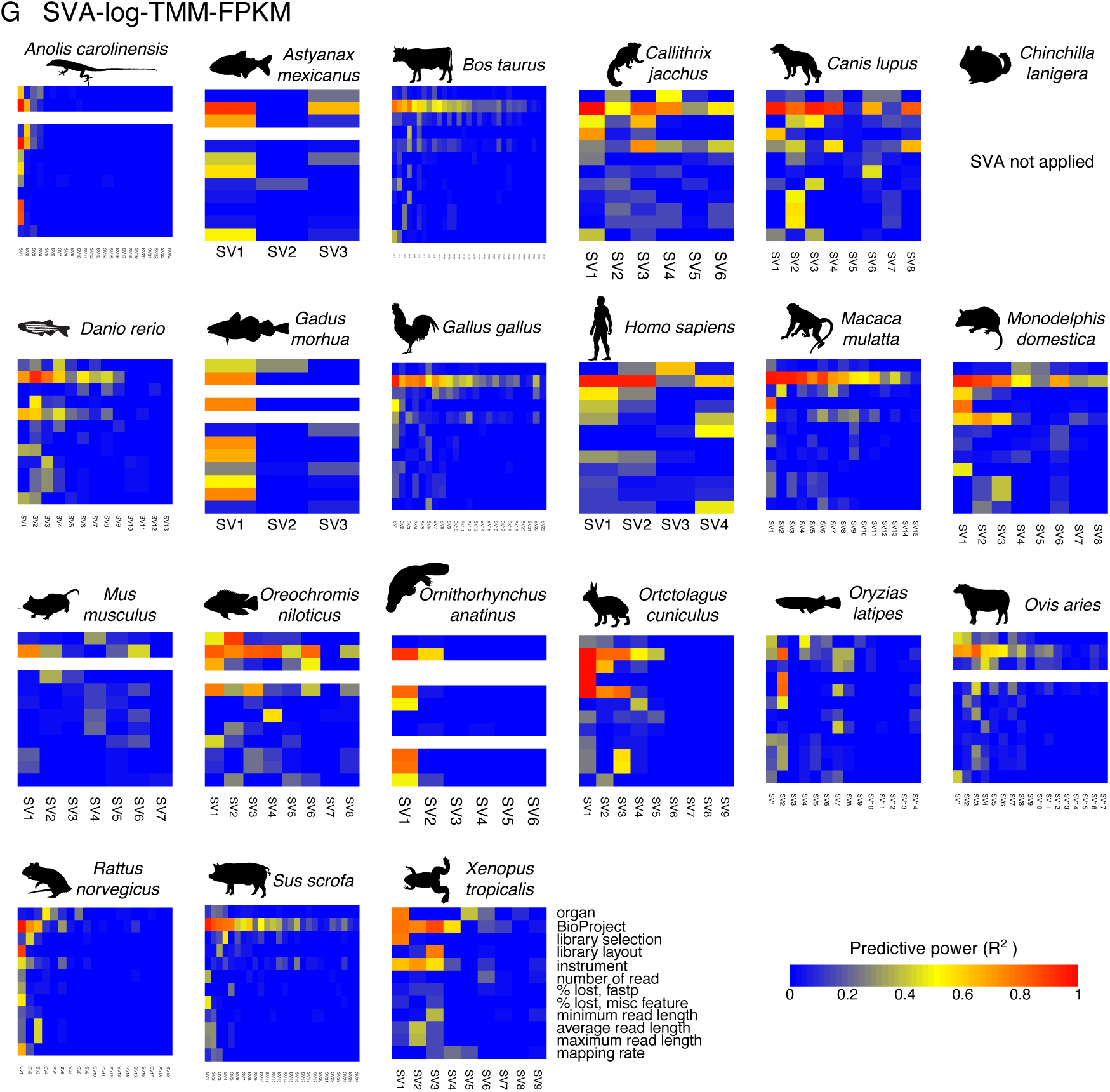
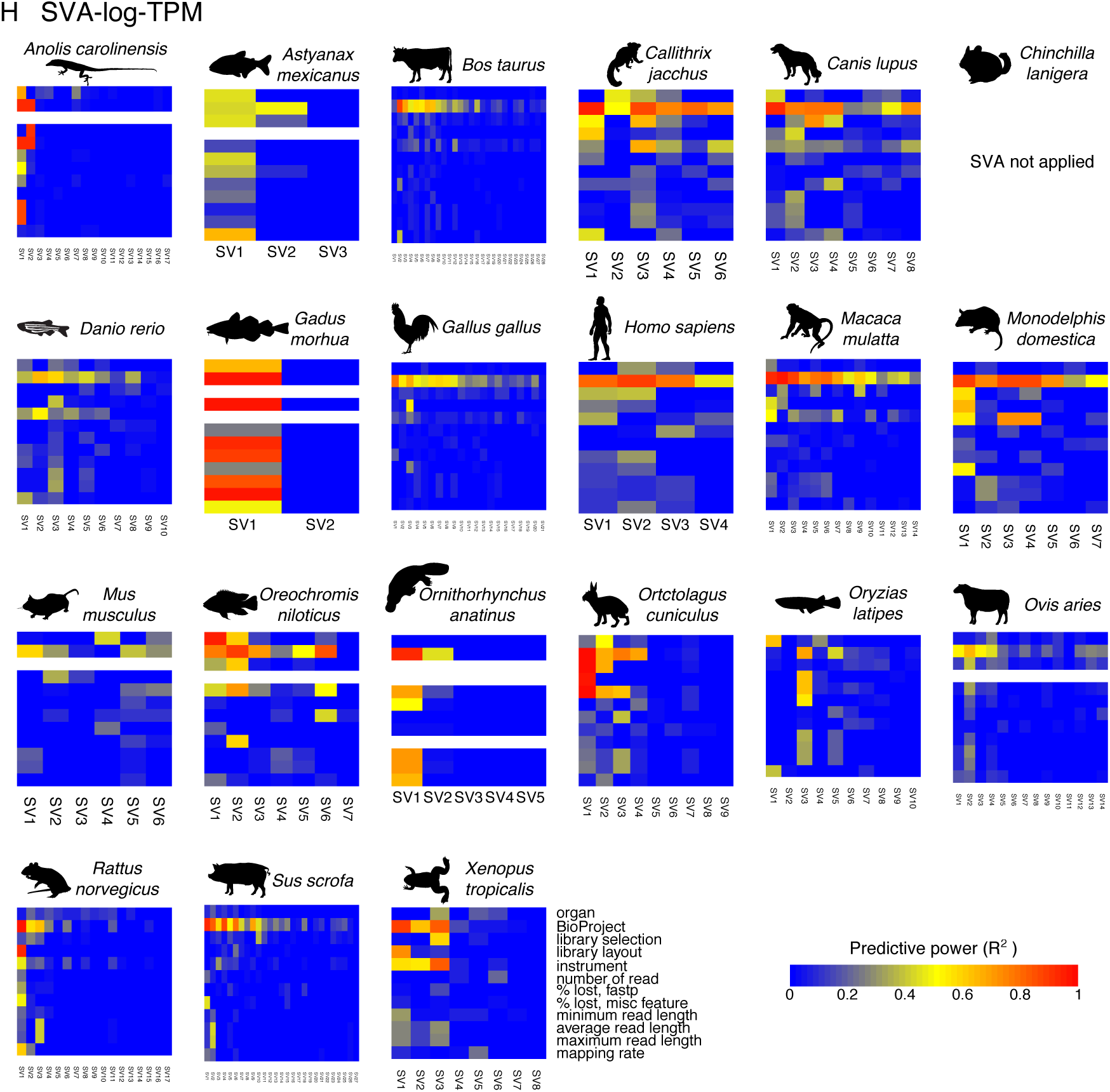
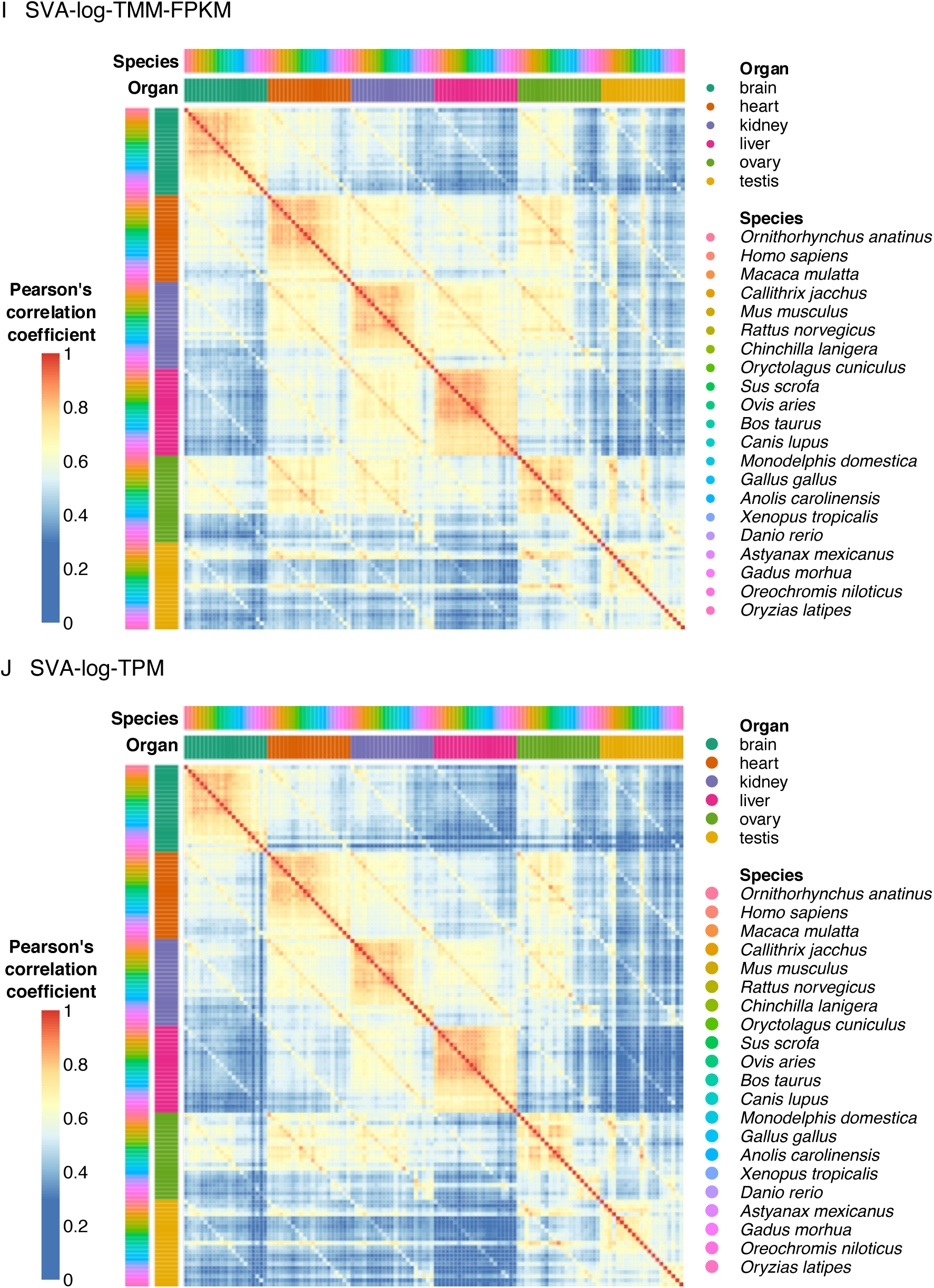
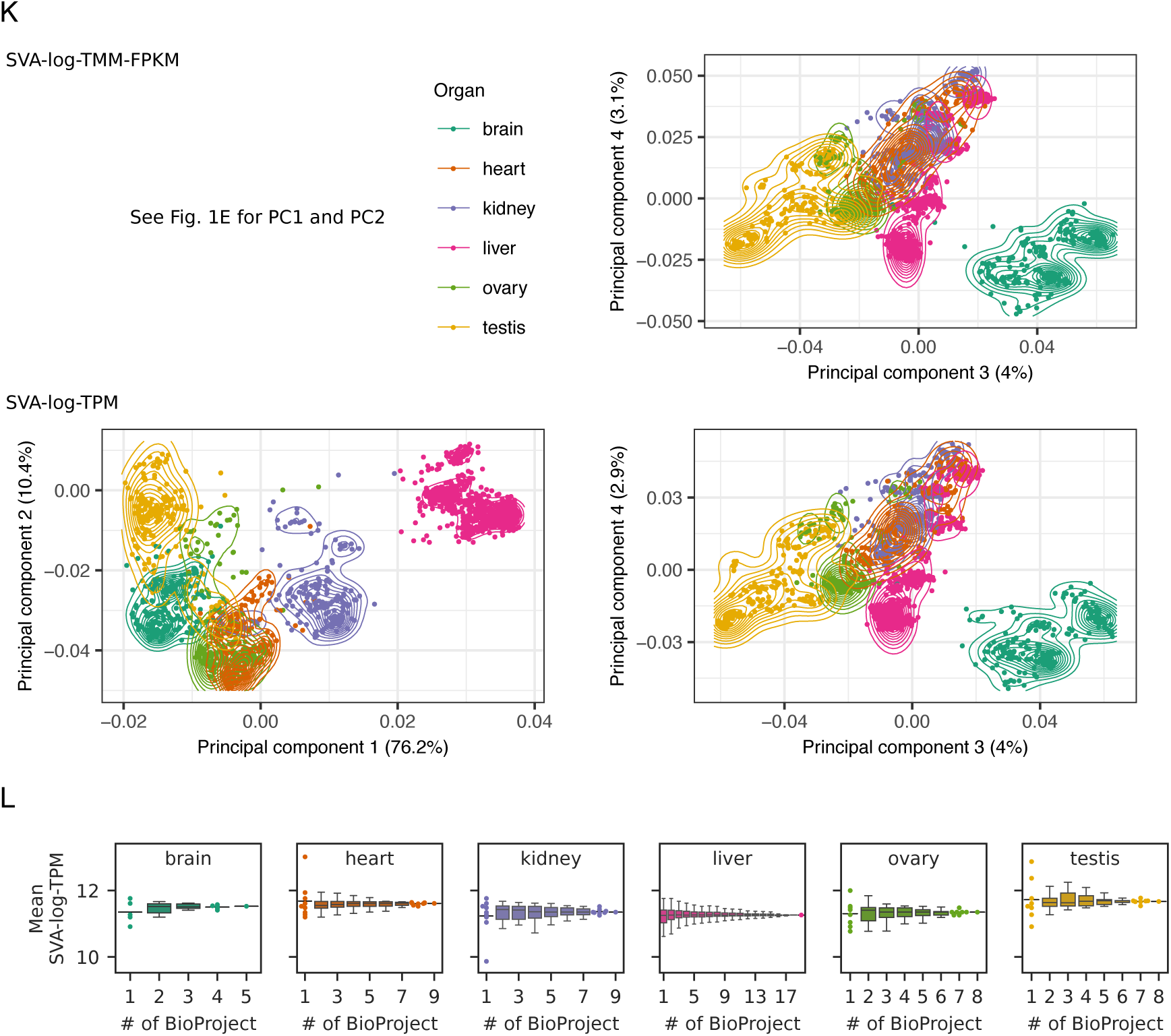
Characteristics of amalgamated transcriptome. **(A)** The number of RNA-seq experiments, BioProjects, and detected surrogate variables in each species. The counts were derived from the final dataset. The numbers of surrogate variables (SV) are correlated with the numbers of RNA-seq samples and BioProjects. (**B**) Relationships of the number of SVs with log-TPM and log-TMM-FPKM. Points correspond to species. (**C–D**) Correlation heatmaps of corrected transcriptomes. See Table S1 for full descriptions including RNA-seq sample IDs and BioProject IDs. (**E–F**) Distinct distributions of Pearson’s correlation coefficients depending on whether a pair of RNA-seq samples are the same organ or whether they are from the same research project. (**G–H**) Predictor analysis of detected surrogate variables. The predictive power was analyzed by linear regression using different properties of RNA-seq experiments: organ (brain, heart, kidney, liver, ovary, and testis), BioProject (e.g., PRJNA176589), library selection (e.g., cDNA and polyA), library layout (single and paired), instrument (e.g., Illumina HiSeq 2500 and NextSeq 550), number of read (e.g., 91,641,467 reads), % lost, fastp (percentage of reads that are removed by fastp; e.g., 5%), % lost, misc feature (percentage of reads that are mapped to non-nuclear-mRNA features and are removed from the analysis; e.g., 5%), minimum read length (e.g., 25 nt), average read length (e.g., 70 nt), maximum read length (e.g., 75 nt), and mapping rate (e.g., 80%). The predictors are summarized in Table S1. (**I–J**) Multispecies correlation analysis of averaged organ expression. Corrected expression levels of 1,377 single-copy orthologs were used to calculate pairwise Pearson’s correlation coefficients. (**K**) A principal component analysis using expression levels of 1,377 single-copy orthologs from 21 species. Points correspond to RNA-seq samples. Curves show the estimated kernel density. Explained variations in percentages are indicated in each axis. (**L**) Estimated organ-wise expression levels of a housekeeping gene. Since data from relatively many BioProjects are available, glyceraldehyde-3-phosphate dehydrogenase gene (GAPDH, ENSGALG00000014442) in *Gallus gallus* is shown. Points correspond to the average expression level calculated by a random subsampling. All data points and the median value (bar), rather than a box plot, are shown if the number of subsampled BioProject combinations is less than 10.

**Fig. S5.**
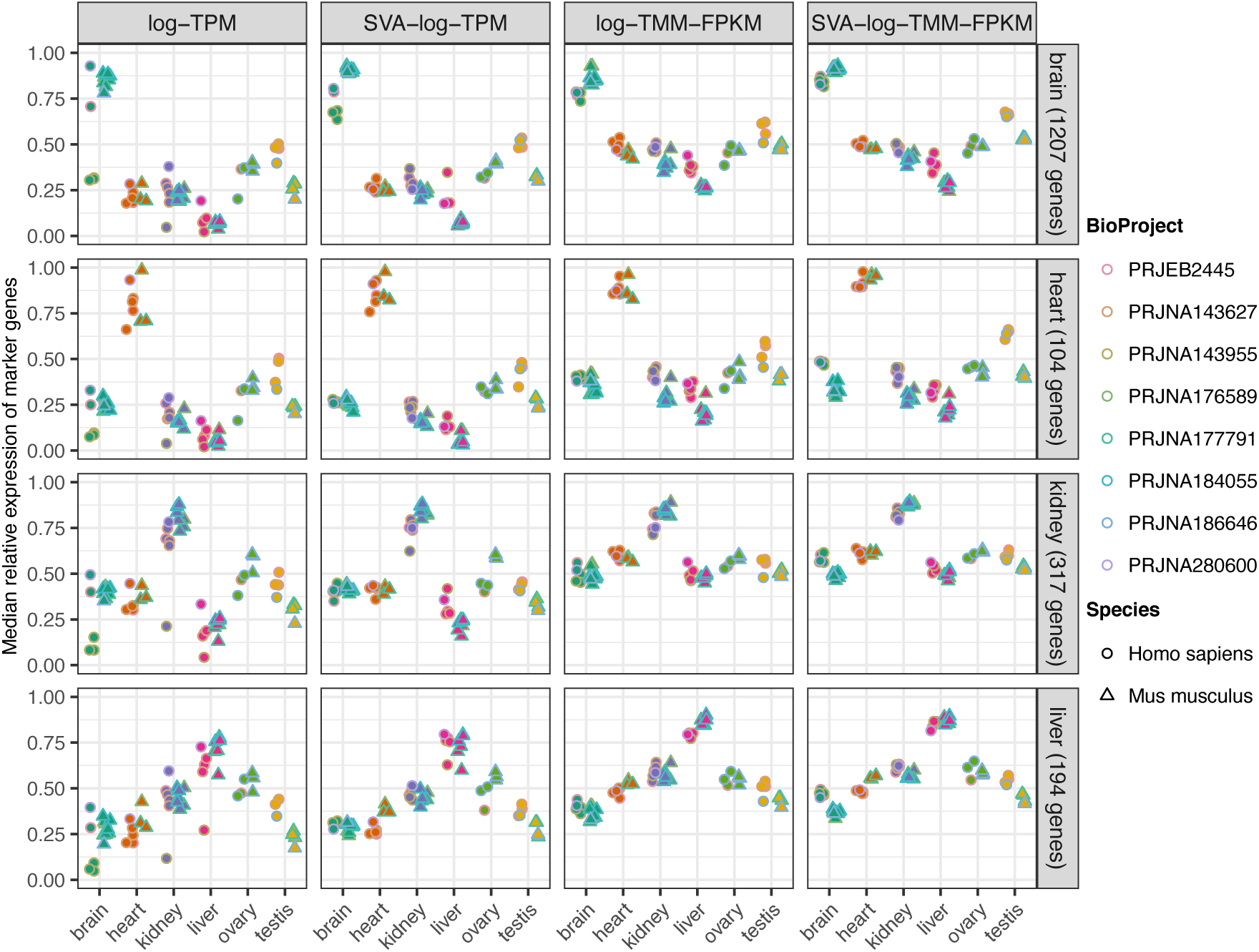
Expression of organ-specific marker genes in human and mouse. Marker genes were retrieved from PanglaoDB (Franzén et al., 2019), and its median expression values were obtained for each RNA-seq sample. A cell-type-wise analysis is provided in Supplementary Data. Cell types in ovary and testis were not included in PanglaoDB (access date: April 1, 2020).

**Fig. S6.**
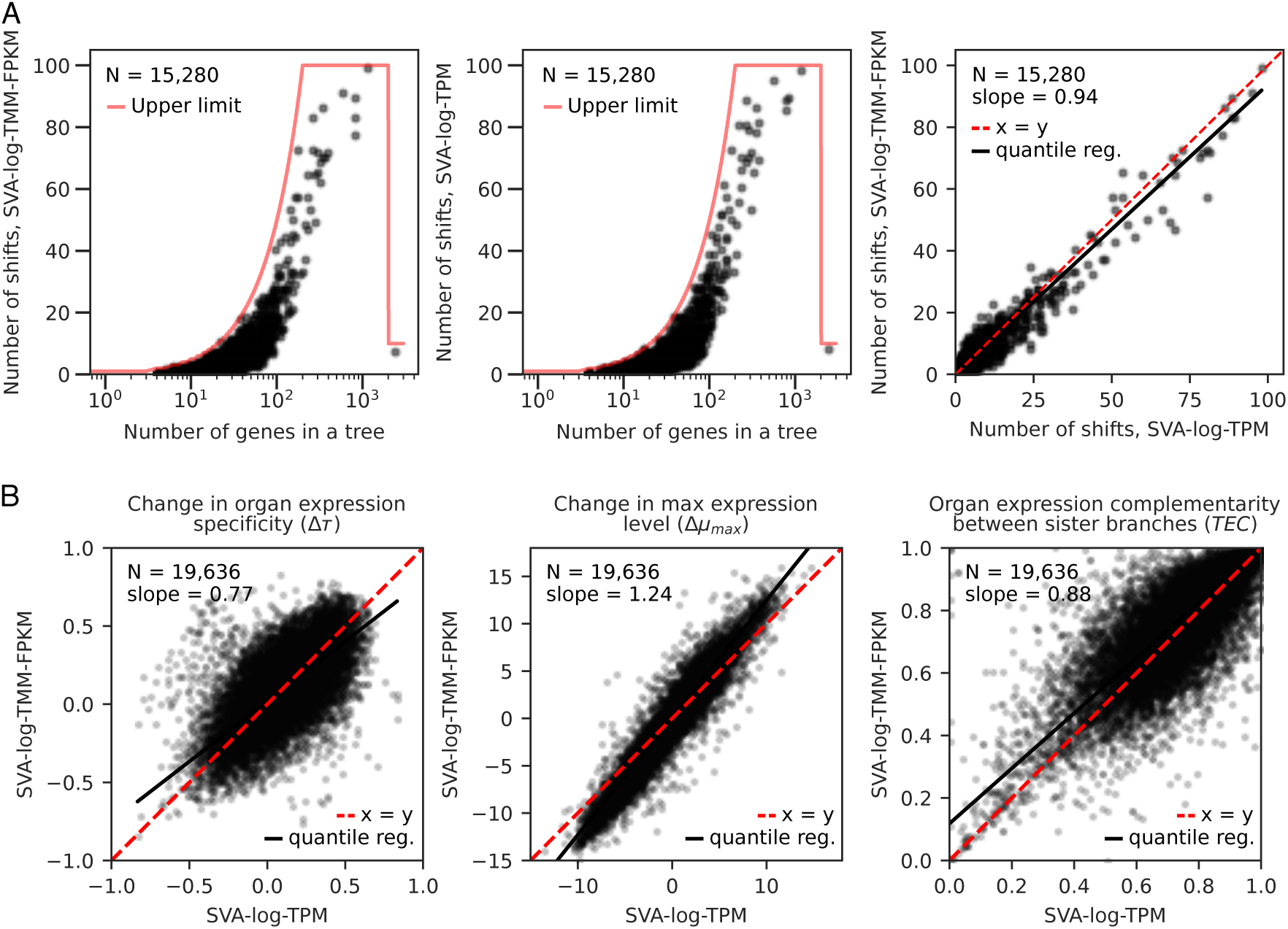
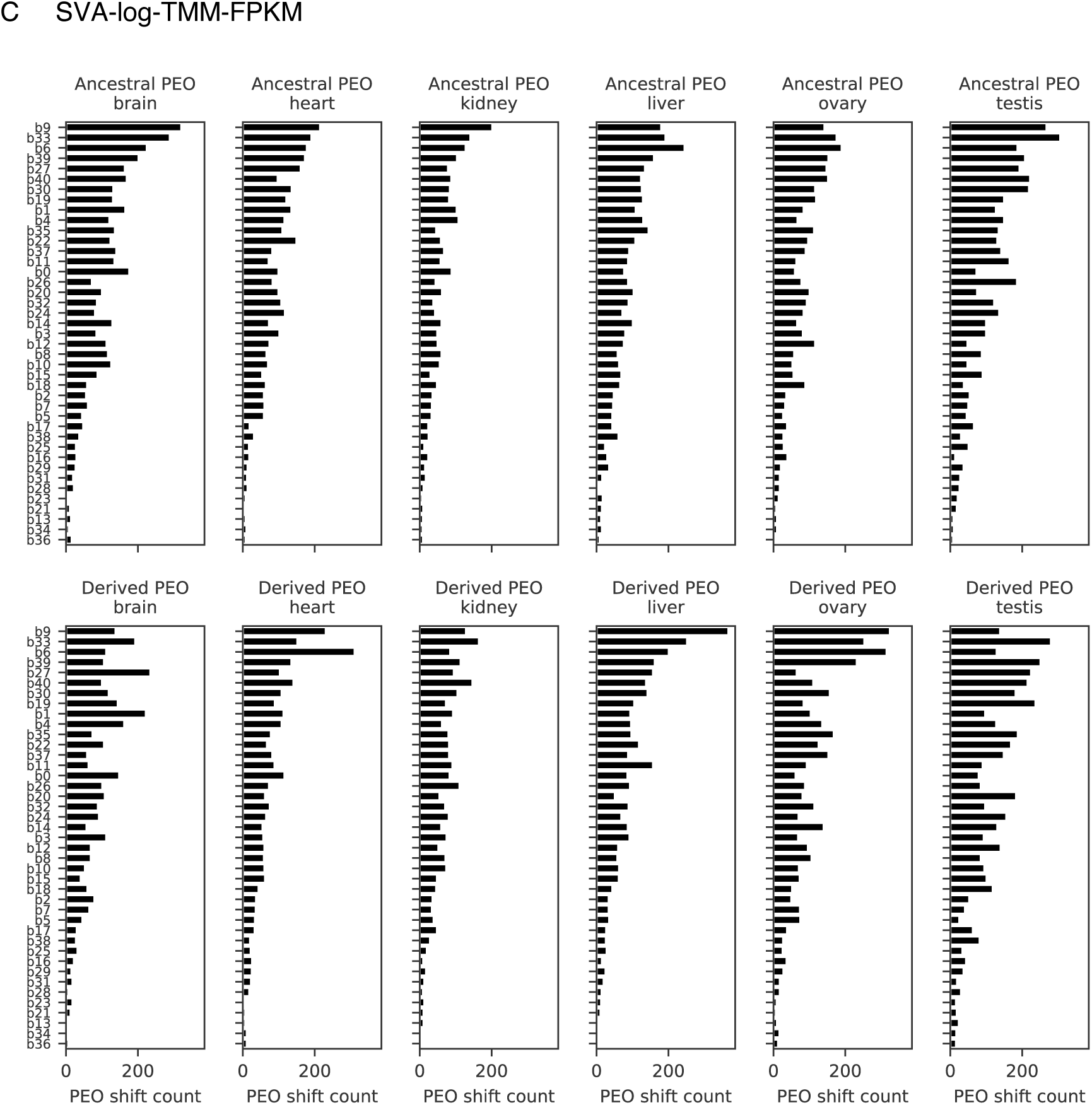
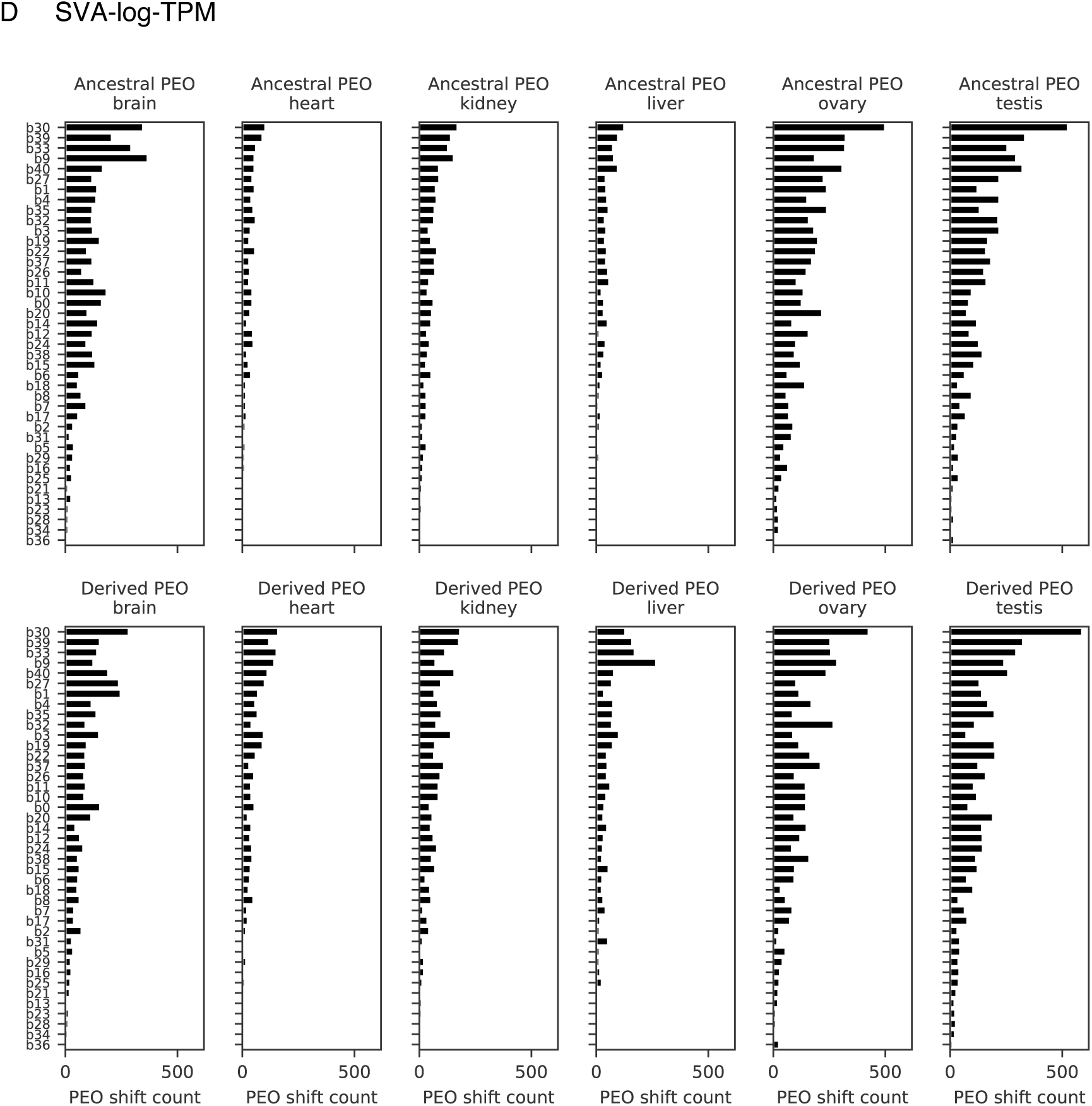
Comparison of SVA-log-FPKM and SVA-log-TPM in expression regime shift detection. (**A**) The numbers of expression regime shifts. Each point corresponds to a gene tree. The solid red lines show the upper limits in the regime shift search. The numbers of shifts detected with SVA-log-TMM-FPKM (left) and SVA-log-TMM (center) are well correlated (right). The black line is a quantile regression, and the dashed red line shows a slope of 1. (**B**) Expression properties. Each point corresponds to an expression regime shift which is consistently detected by SVA-log-TMM-FPKM and SVA-log-TPM. (**C–D**) The branch-wise numbers of detected shifts in the species tree. The shifts were categorized by primary-expressed organs (PEOs) in ancestral and derived states. The order of the species tree branches (y axis) corresponds to the total number of corresponding gene tree branches in the dataset. See Fig. 3A for branch ids.

**Fig. S7.**
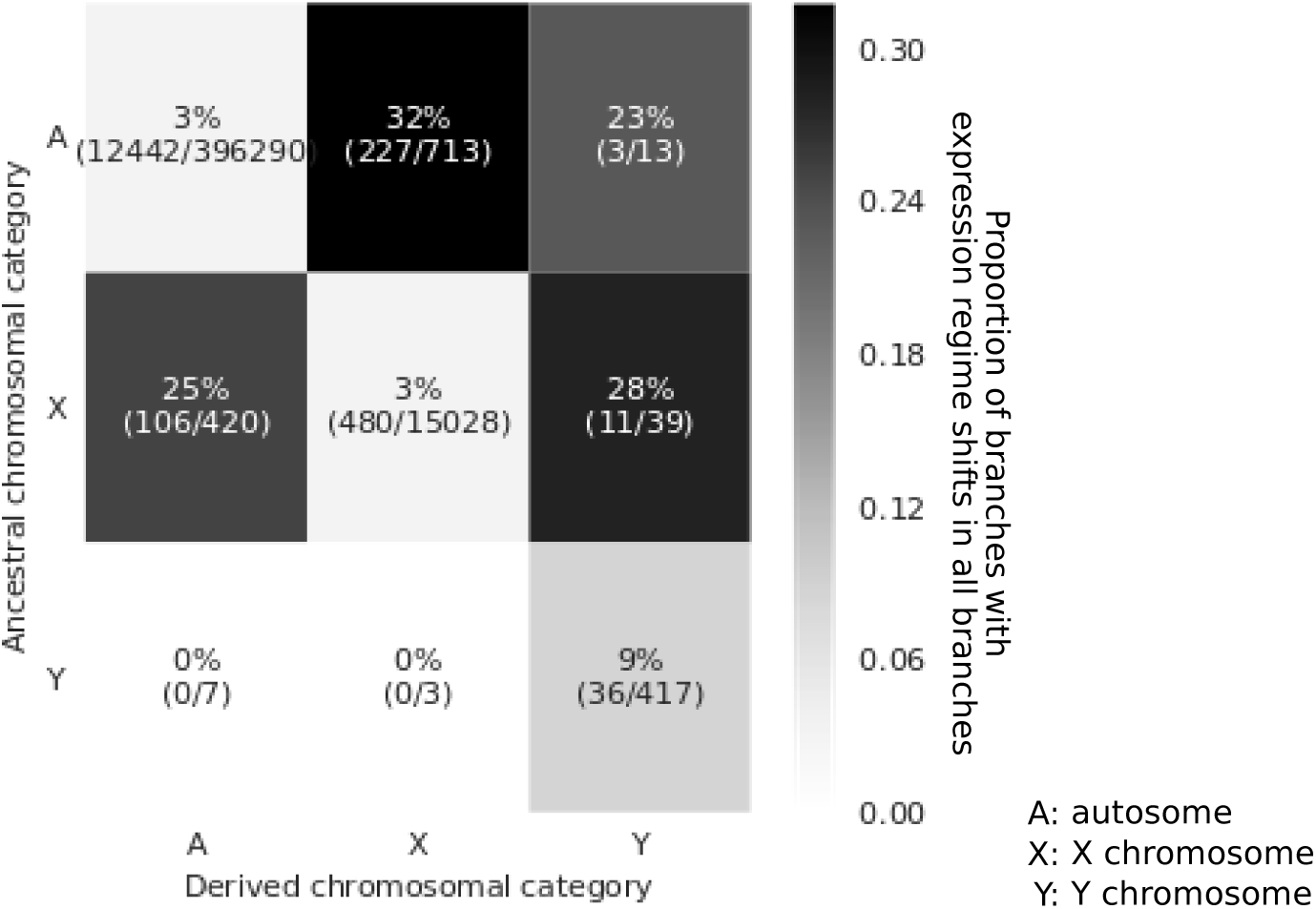
The relationships between expression shifts and chromosomal location. The heatmap shows the frequency of expression shifts observed among the branches with or without a change in the chromosomal category (non-diagonal or diagonal, respectively). Chromosomal locations were categorized into autosomes (A), X chromosome (X) and Y chromosome (Y), and the ancestral locations were inferred by stochastic character mapping.

**Fig. S8.**
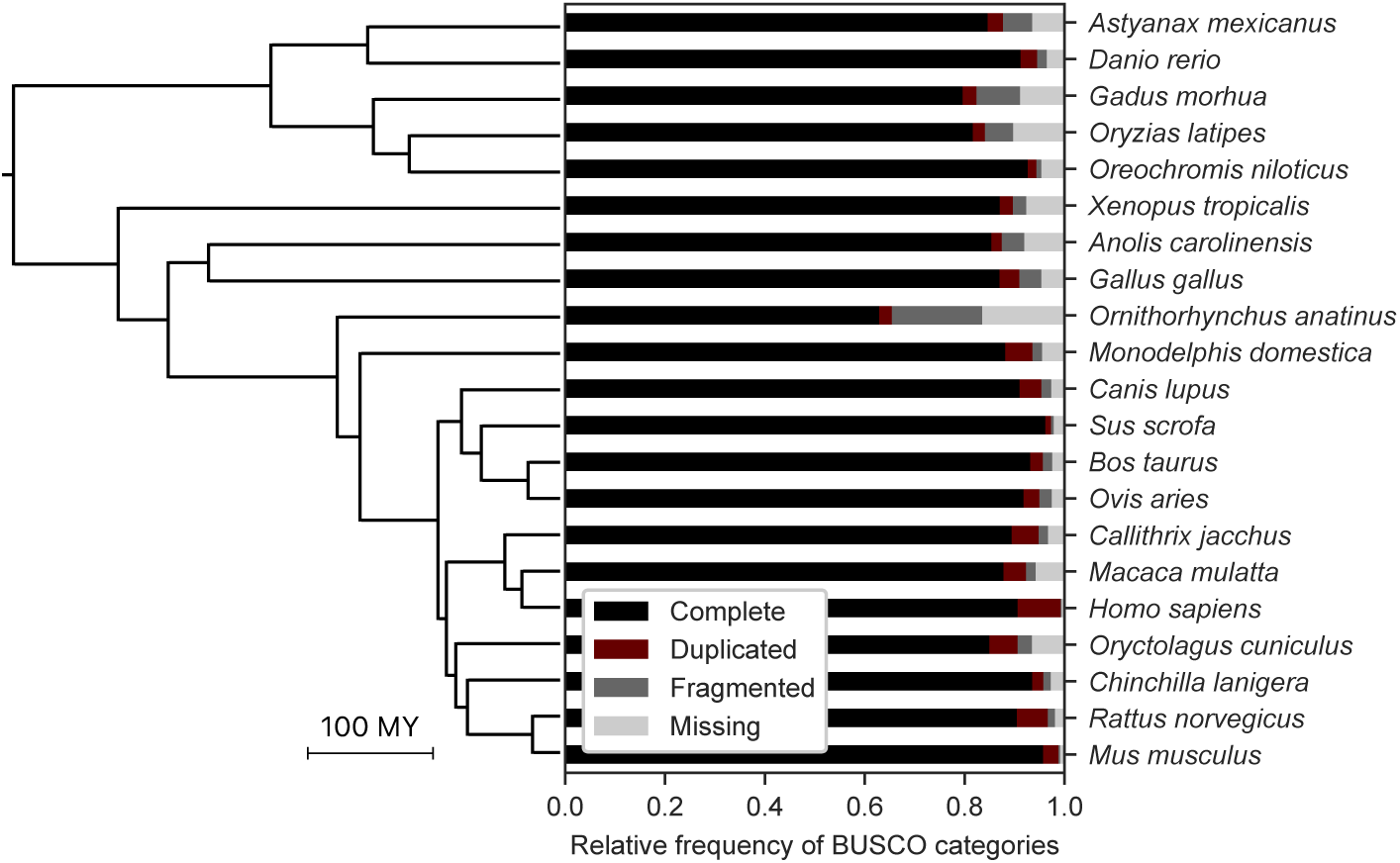
Evaluation of gene set completeness. BUSCO analysis was performed for gene sets from 21 species using 3,407 single-copy orthologs in the dataset “vertebrata_odb10”. The species tree is shown to visualize lineage-specific trends.

**Fig. S9.**
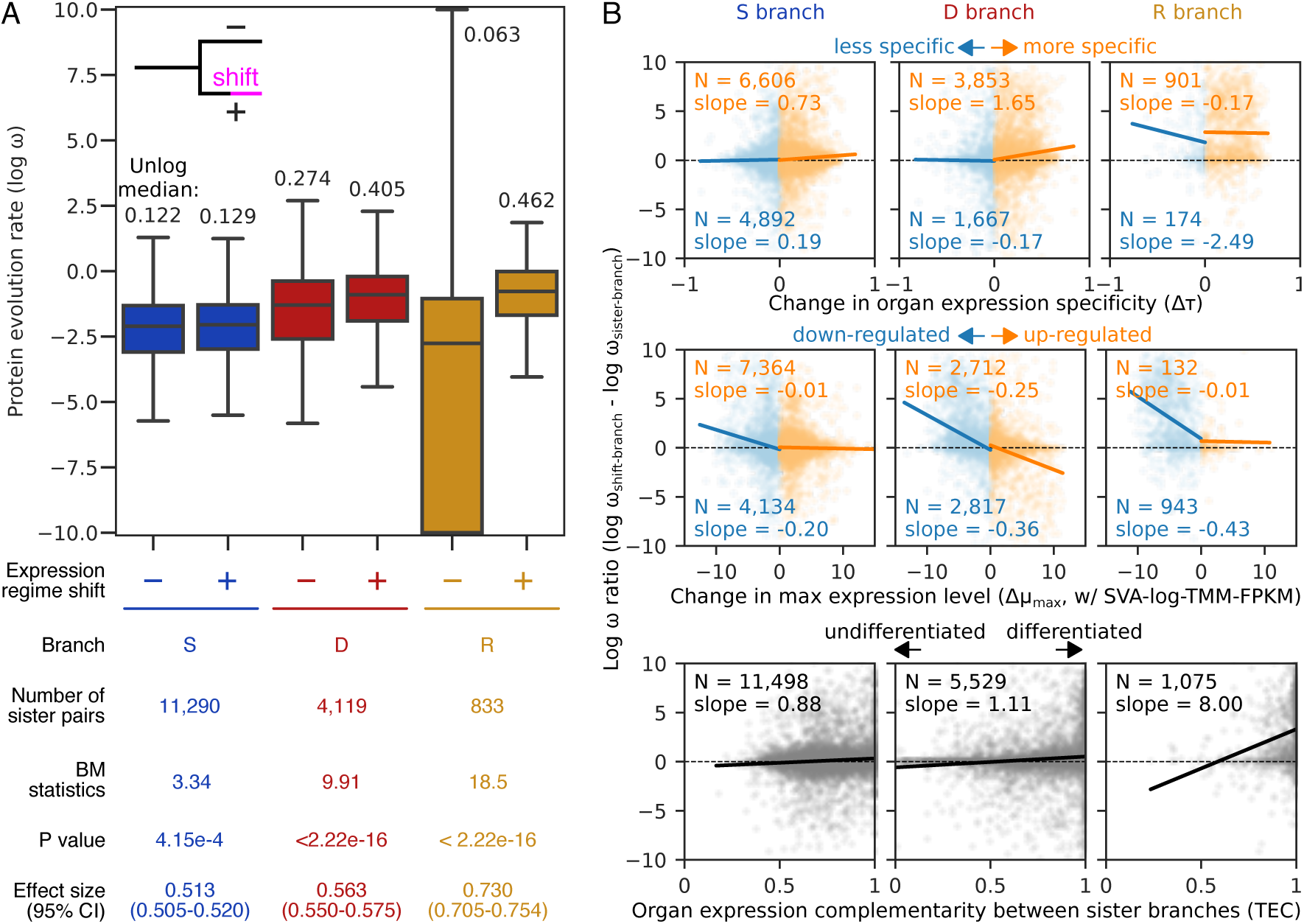
Alternative analysis supporting non-linear change in protein evolution rate in correlation with expression regime shifts. (**A**) Distribution of ω values. While the results from stochastic character mapping (map*dNdS*) are reported in the main text, results from maximum-likelihood estimation (HyPhy) are shown here. A plus (+) indicates branches with expression shifts, whereas minus (−) branches are sisters to the ‘plus’ branches. Statistical differences between pairs of distributions were tested using a two-sided Brunner–Munzel test (Brunner and Munzel, 2000). Non-log-transformed median values are shown above the boxplots. For visualization purpose, extreme values exceeding ±10 were clipped. (**B**) Relationships between protein evolution rate and change in expression properties. While SVA-log-TMM-FPKM-based analysis is reported in the main text, results from SVA-lot-TPM-based analysis is shown here. Stochastic character mapping was used to obtain branch-wise ω values. Points correspond to expression regime shifts (log ω ratio = 0). Dashed lines indicate no between-branch difference in protein evolution rate. Solid lines show a linear regression. Its slope and number of regime shifts are also provided. Regime shifts with negative and positive changes were separately analyzed for organ specificity (upper) and expression level (middle).

**Fig. S10.**
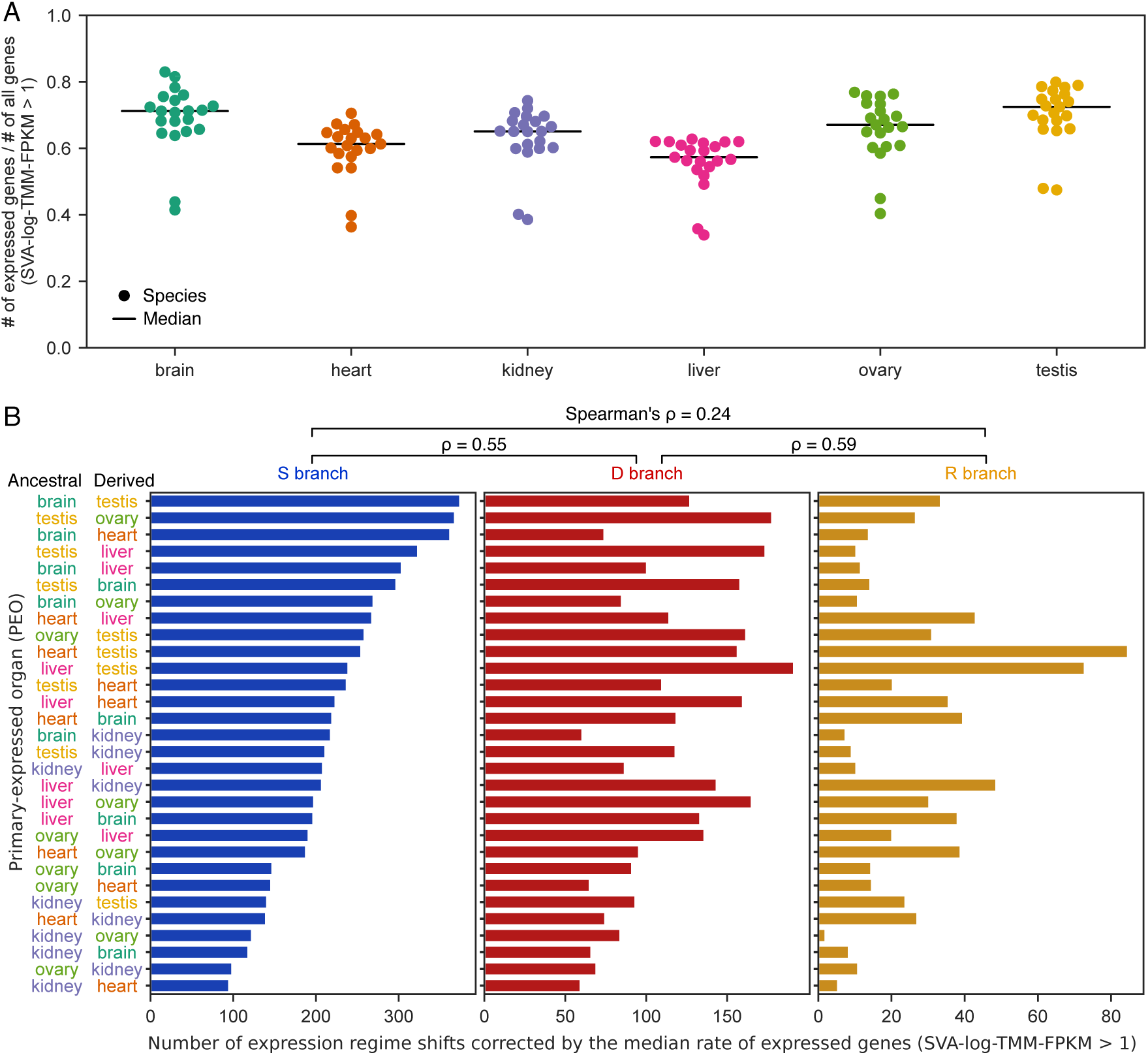
The number of expressed genes does not explain the organ-wise abundance of PEO shifts. (**A**) The organ-wise numbers of expressed genes in the 21 species. In this analysis, expressed genes are defined as genes with >1 SVA-log-TMM-FPKM. (**B**) The PEO shift distributions corrected by the median numbers of expressed genes. The data in Fig. 5A were corrected as follows: 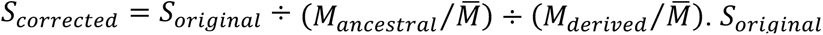 corresponds to the numbers shown in Fig. 5A. *M_ancestral_* and *M_derived_* are median numbers of expressed genes in ancestral and derived PEOs, respectively. 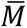 indicates the across-organ average of expressed gene numbers.

**Fig. S11.**
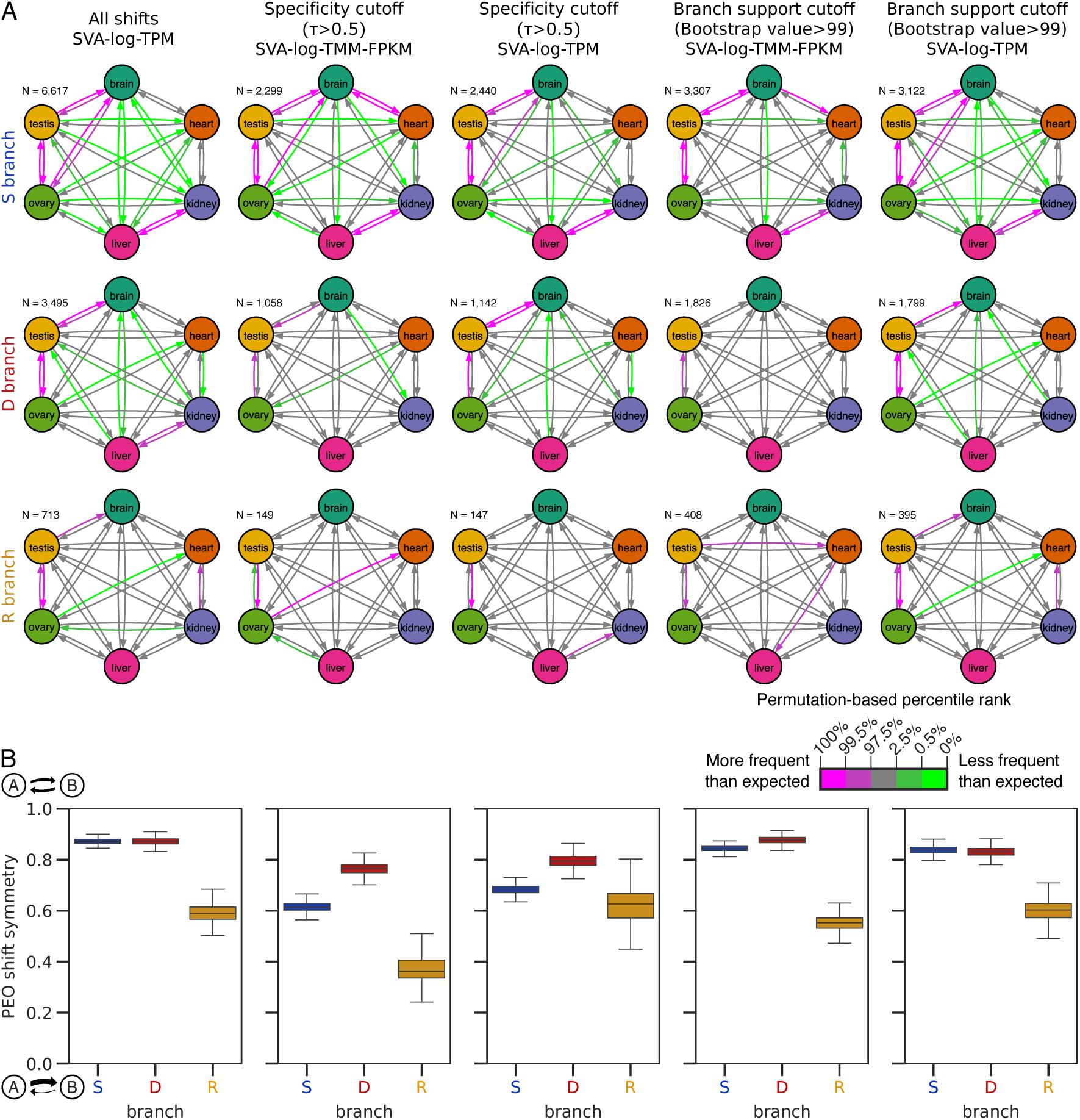
Evolutionary dynamics of gene expression analyzed with conservative datasets. Both SVA-log-TPM and SVA-log-TMM FPKM values were analyzed. Organ-specific genes were examined by selecting PEO shifts with high ancestral and derived organ expression specificity (τ > 0.5). To examine the effect of inference errors in gene tree reconstruction, branches with high support values (ultrafast bootstrap percentage > 99) were analyzed. **A** and **B** correspond to Fig. 5B and C, respectively.

**Fig. S12.**
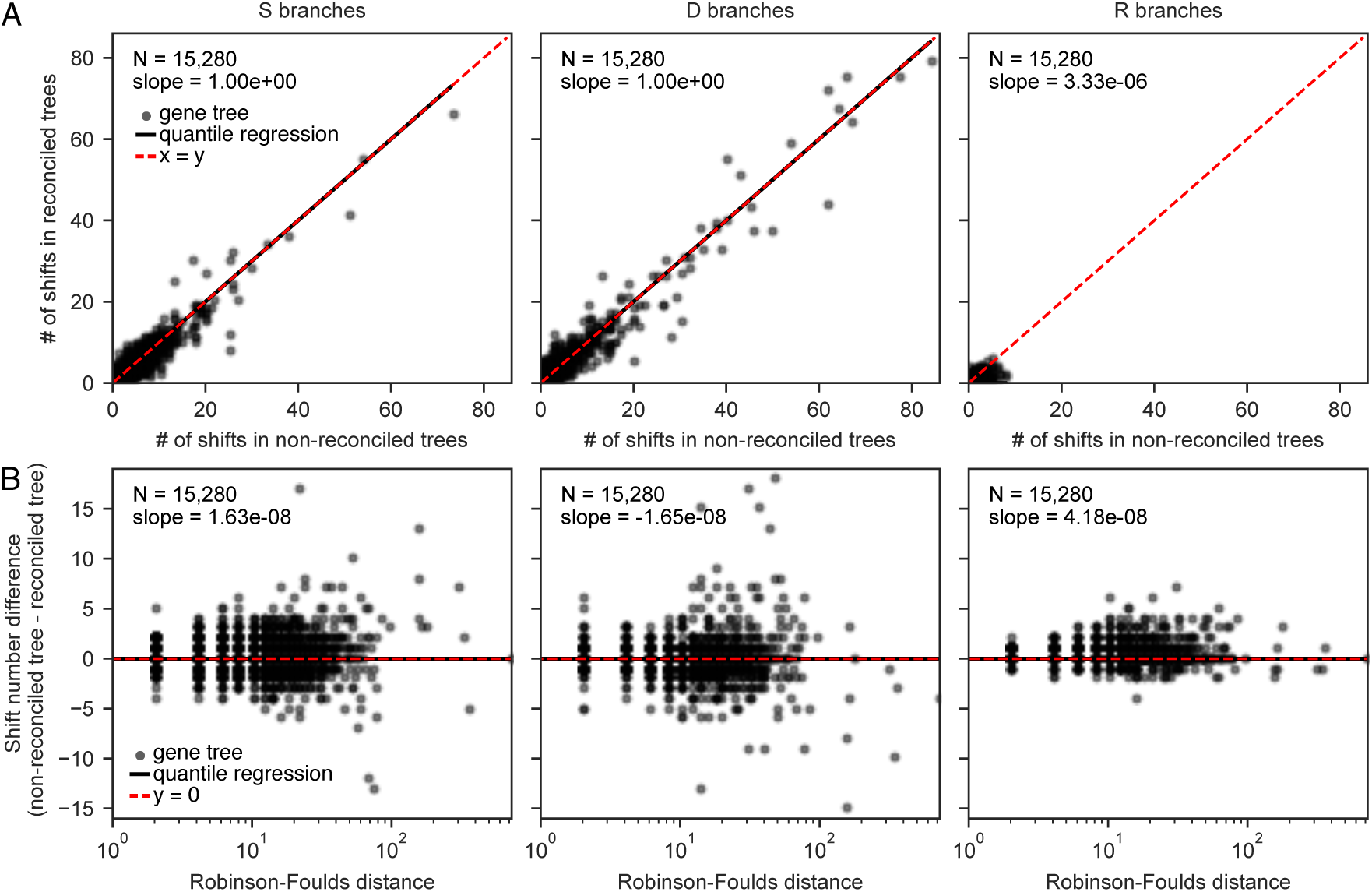
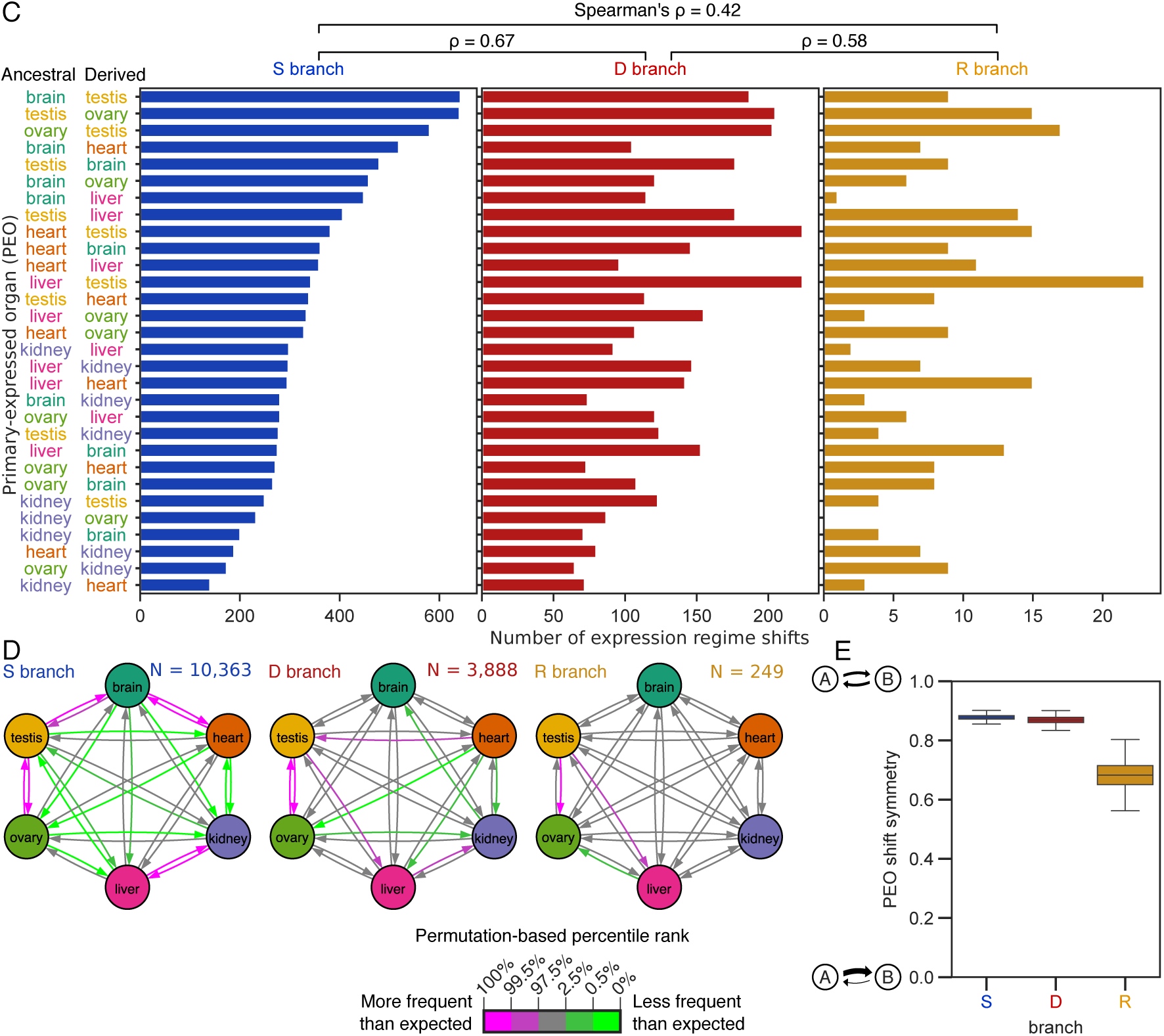
Effects of phylogeny reconciliation. (**A**) The numbers of detected shifts in non-reconciled and reconciled maximum-likelihood gene trees. (**B**) The relationship between shift number difference and tree topology difference measured by Robinson-Foulds distance ^51^. Trees with no detected shifts were removed from the analysis. (**C–E**) Reproduction of PEO shift distributions using reconciled gene trees. C, D, and E correspond to Fig. 5A, Fig. 5B, and Fig. 5C, respectively.

**Fig. S13.**
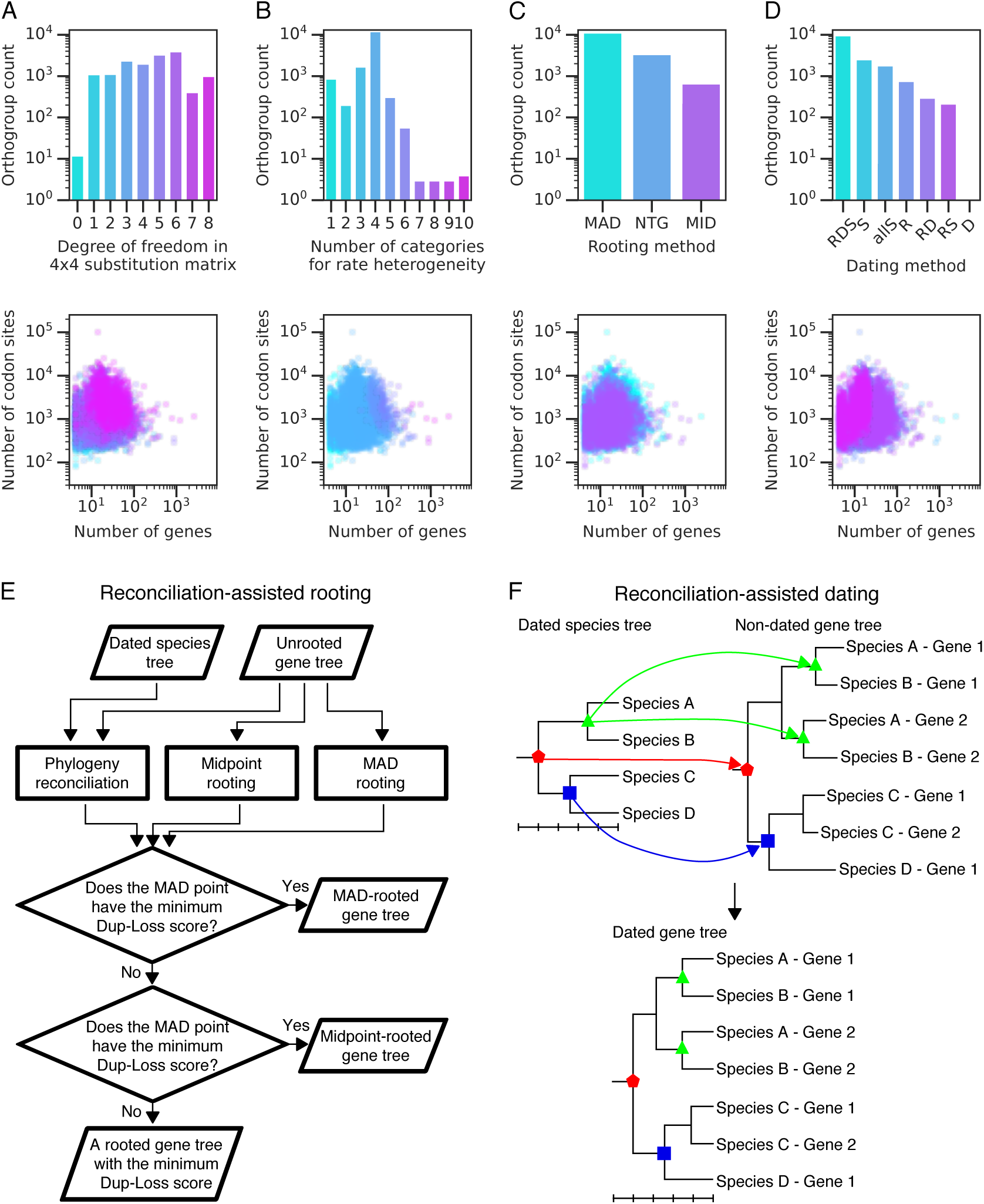
Gene tree reconstruction. (**A**) Complexity of best-fit nucleotide substitution matrices. (**B**) Complexity of rate heterogeneity among nucleotide sites. Both discrete Gamma models (Yang, 1994) and FreeRate models (Soubrier et al., 2012; Yang, 1995) were included to count the number of categories for rate heterogeneity. (**C**) Selected rooting positions in reconciliation-assisted gene tree rooting. MAD, minimal ancestor deviation; NTG, ‘rooting mode’ of NOTUNG; MID, midpoint between the longest path. (**D**) Time-constrained nodes in tree dating. R, root node; S, speciation node; D, duplication node. All available constraints (RDS) are used in the first trial and then successively relaxed if estimation fails. In the category ‘allS’, all nodes are speciation nodes and therefore no divergence time estimation was performed for the trees. (**E**) Reconciliation-assisted gene tree rooting. Rooting points were estimated with two different methods: the minimum ancestor deviation (MAD) method and midpoint rooting. If they were compatible with the event parsimony involving gene duplication and loss, the MAD- or midpoint-rooted tree was reported in sequence. If not, one of event parsimony trees was reported. (**F**) Reconciliation-assisted gene tree dating. Speciation nodes in the dated species tree were mapped onto the non-dated gene tree by phylogeny reconciliation. By using those nodes as calibration points, the other node ages were estimated using the penalized likelihood method.

**Fig. S14.**
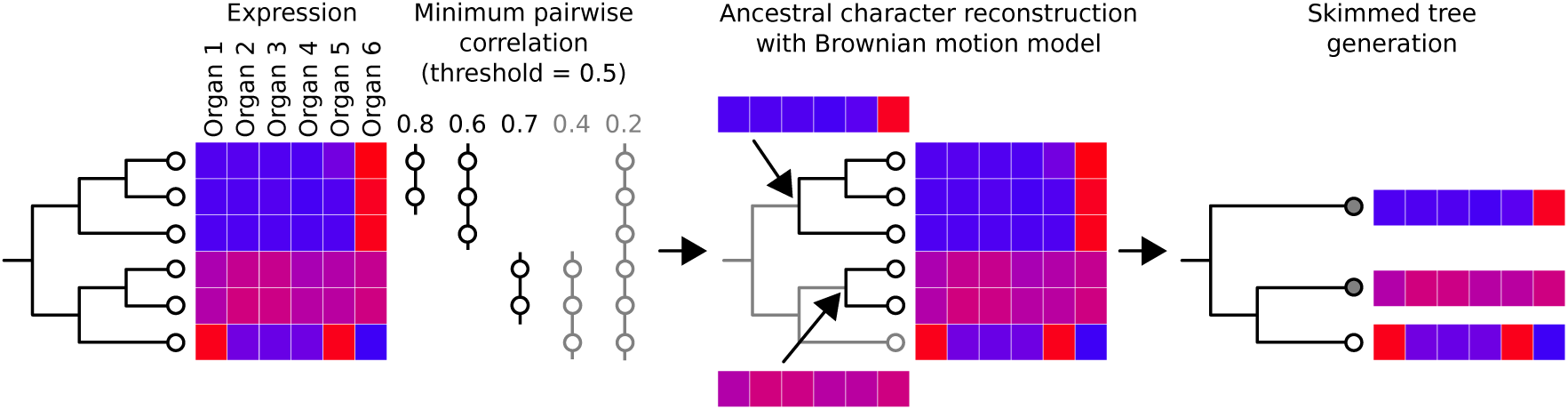
Gene tree skimming in phylogenetic comparative analysis. Gene tree clades are collapsed if character states are highly correlated. Resultant trees contain a smaller number of leaves than the original trees while preserving drastic changes in character evolution. Note that, unlike this example, we used extremely stringent threshold in the analysis (Pearson’s correlation coefficient > 0.99).

## References

1. Altenhoff, A.M., Studer, R.A., Robinson-Rechavi, M., Dessimoz, C., and Couto, F. (2012). Resolving the ortholog conjecture: orthologs tend to be weakly, but significantly, more similar in function than paralogs. PLoS Comput. Biol. 8, e1002514.

2. Assis, R., and Bachtrog, D. (2015). Rapid divergence and diversification of mammalian duplicate gene functions. BMC Evol. Biol. 15, 138.

3. Babushok, D. V., Ostertag, E.M., and Kazazian, H.H. (2007). Current topics in genome evolution: Molecular mechanisms of new gene formation. Cell. Mol. Life Sci. 64, 542– 554.

4. Balakirev, E.S., and Ayala, F.J. (2003). Pseudogenes: Are they “junk” or functional DNA? Annu. Rev. Genet. 37, 123–151.

5. Barbosa-Morais, N.L., Irimia, M., Pan, Q., Xiong, H.Y., Gueroussov, S., Lee, L.J., Slobodeniuc, V., Kutter, C., Watt, S., Colak, R., et al. (2012). The evolutionary landscape of alternative splicing in vertebrate species. Science 338, 1587–1593.

6. Bedford, T., and Hartl, D.L. (2009). Optimization of gene expression by natural selection. Proc. Natl. Acad. Sci. U. S. A. 106, 1133–1138.

7. Boer, P.H., Adra, C.N., Lau, Y.F., and McBurney, M.W. (1987). The testis-specific phosphoglycerate kinase gene pgk-2 is a recruited retroposon. Mol. Cell. Biol. 7, 3107– 3112.

8. Bollback, J. (2006). SIMMAP: Stochastic character mapping of discrete traits on phylogenies. BMC Bioinformatics 7, 88.

9. Brawand, D., Soumillon, M., Necsulea, A., Julien, P., Csárdi, G., Harrigan, P., Weier, M., Liechti, A., Aximu-Petri, A., Kircher, M., et al. (2011). The evolution of gene expression levels in mammalian organs. Nature 478, 343–348.

10. Bray, N.L., Pimentel, H., Melsted, P., and Pachter, L. (2016). Near-optimal probabilistic RNA-seq quantification. Nat. Biotechnol. 34, 525–527.

11. Breschi, A., Djebali, S., Gillis, J., Pervouchine, D.D., Dobin, A., Davis, C.A., Gingeras, T.R., and Guigó, R. (2016). Gene-specific patterns of expression variation across organs and species. Genome Biol. 17, 151.

12. Brunner, E., and Munzel, U. (2000). The nonparametric Behrens-Fisher problem: asymptotic theory and a small-sample approximation. Biom. J. 42, 17–25.

13. Budd, G.E. (2007). On the origin and evolution of major morphological characters. Biol. Rev. 81, 609–628.

14. Capella-Gutierrez, S., Silla-Martinez, J.M., and Gabaldon, T. (2009). trimAl: a tool for automated alignment trimming in large-scale phylogenetic analyses. Bioinformatics 25, 1972–1973.

15. Cardoso-Moreira, M., Halbert, J., Valloton, D., Velten, B., Chen, C., Shao, Y., Liechti, A., Ascenção, K., Rummel, C., Ovchinnikova, S., et al. (2019). Gene expression across mammalian organ development. Nature 571, 505–509.

16. Carelli, F.N., Hayakawa, T., Go, Y., Imai, H., Warnefors, M., and Kaessmann, H. (2016). The life history of retrocopies illuminates the evolution of new mammalian genes. Genome Res. 26, 301–314.

17. Castillo-Davis, C.I., Hartl, D.L., and Achaz, G. (2004). cis-Regulatory and protein evolution in orthologous and duplicate genes. Genome Res. 14, 1530–1536.

18. Chen, X., and Zhang, J. (2012). The ortholog conjecture is untestable by the current gene ontology but is supported by RNA sequencing data. PLoS Comput. Biol. 8, e1002784.

19. Chen, J., Swofford, R., Johnson, J., Cummings, B.B., Rogel, N., Lindblad-Toh, K., Haerty, W., Palma, F. di, and Regev, A. (2019). A quantitative framework for characterizing the evolutionary history of mammalian gene expression. Genome Res. 29, 53–63.

20. Chen, K., Durand, D., and Farach-Colton, M. (2000). NOTUNG: a program for dating gene duplications and optimizing gene family trees. J. Comput. Biol. 7, 429–447.

21. Chen, S., Krinsky, B.H., and Long, M. (2013). New genes as drivers of phenotypic evolution. Nat. Rev. Genet. 14, 645–660.

22. Chen, S., Zhou, Y., Chen, Y., and Gu, J. (2018). fastp: an ultra-fast all-in-one FASTQ preprocessor. Bioinformatics 34, i884–i890.

23. Conant, G.C., and Wolfe, K.H. (2008). Turning a hobby into a job: How duplicated genes find new functions. Nat. Rev. Genet. 9, 938–950.

24. Cortez, D., Marin, R., Toledo-Flores, D., Froidevaux, L., Liechti, A., Waters, P.D., Grützner, F., and Kaessmann, H. (2014). Origins and functional evolution of Y chromosomes across mammals. Nature 508, 488–493.

25. Danshina, P. V., Geyer, C.B., Dai, Q., Goulding, E.H., Willis, W.D., Kitto, G.B., McCarrey, J.R., Eddy, E.M., and O’Brien, D.A. (2010). Phosphoglycerate kinase 2 (PGK2) is essential for sperm function and male fertility in mice. Biol. Reprod. 82, 136–145.

26. de-Leon, S.B.-T., and Davidson, E.H. (2007). Gene regulation: gene control network in development. Annu. Rev. Biophys. Biomol. Struct. 36, 191–212.

27. Des Marais, D.L., and Rausher, M.D. (2008). Escape from adaptive conflict after duplication in an anthocyanin pathway gene. Nature 454, 762–765.

28. Dong, D., Yuan, Z., and Zhang, Z. (2011). Evidences for increased expression variation of duplicate genes in budding yeast: from *cis*- to *trans*- regulation effects. Nucleic Acids Res. 39, 837–847.

29. Emms, D.M., and Kelly, S. (2015). OrthoFinder: solving fundamental biases in whole genome comparisons dramatically improves orthogroup inference accuracy. Genome Biol. 16, 157.

30. Farris, J.S. (1972). Estimating phylogenetic trees from distance matrices. Am. Nat. 106, 645–668.

31. Franzén, O., Gan, L.-M., and Björkegren, J.L.M. (2019). PanglaoDB: a web server for exploration of mouse and human single-cell RNA sequencing data. Database 2019.

32. Gessi, M., Hammes, J., Lauriola, L., Dörner, E., Kirfel, J., Kristiansen, G., Muehlen, A. zur, Denkhaus, D., Waha, A., and Pietsch, T. (2013). GNA11 and N-RAS mutations: alternatives for MAPK pathway activating GNAQ mutations in primary melanocytic tumours of the central nervous system. Neuropathol. Appl. Neurobiol. 39, 417–425.

33. Gould, S.J., and Vrba, E.S. (1982). Exaptation—a missing term in the science of form. Paleobiology 8, 4–15.

34. Gouveia-Oliveira, R., Sackett, P.W., and Pedersen, A.G. (2007). MaxAlign: maximizing usable data in an alignment. BMC Bioinformatics 8, 312.

35. Guéguen, L., and Duret, L. (2018). Unbiased estimate of synonymous and nonsynonymous substitution rates with nonstationary base composition. Mol. Biol. Evol. 35, 734–742.

36. Guéguen, L., Gaillard, S., Boussau, B., Gouy, M., Groussin, M., Rochette, N.C., Bigot, T., Fournier, D., Pouyet, F., Cahais, V., et al. (2013). Bio++: efficient extensible libraries and tools for computational molecular evolution. Mol. Biol. Evol. 30, 1745– 1750.

37. Guschanski, K., Warnefors, M., and Kaessmann, H. (2017). The evolution of duplicate gene expression in mammalian organs. Genome Res. 27, 1461–1474.

38. Hansen, T.F. (1997). Stabilizing selection and the comparative analysis of adaptation. Evolution 51, 1341–1351.

39. Hedges, S.B., Dudley, J., and Kumar, S. (2006). TimeTree: a public knowledge-base of divergence times among organisms. Bioinformatics 22, 2971–2972.

40. Hoang, D.T., Chernomor, O., von Haeseler, A., Minh, B.Q., and Vinh, L.S. (2018). UFBoot2: Improving the ultrafast bootstrap approximation. Mol. Biol. Evol. 35, 518– 522.

41. Huerta-Cepas, J., Dopazo, H., Dopazo, J., Gabaldón, T., McPherson, J., Marra, M., Hillier, L., Waterston, R., Chinwalla, A., Wallis, J., et al. (2007). The human phylome. Genome Biol. 8, 934–941.

42. Huerta-Cepas, J., Dopazo, J., Huynen, M.A., and Gabaldón, T. (2011). Evidence for short-time divergence and long-time conservation of tissue-specific expression after gene duplication. Brief. Bioinform. 12, 442–448.

43. Huerta-Cepas, J., Serra, F., and Bork, P. (2016). ETE 3: Reconstruction, analysis, and visualization of phylogenomic data. Mol. Biol. Evol. 33, 1635–1638.

44. Julien, P., Brawand, D., Soumillon, M., Necsulea, A., Liechti, A., Schütz, F., Daish, T., Grützner, F., and Kaessmann, H. (2012). Mechanisms and evolutionary patterns of mammalian and avian dosage compensation. PLoS Biol. 10, e1001328.

45. Jun, J., Ryvkin, P., Hemphill, E., Mandoiu, I., and Nelson, C. (2009). The birth of new genes by RNA- and DNA-mediated duplication during mammalian evolution. J. Comput. Biol. 16, 1429–1444.

46. Kaessmann, H. (2010). Origins, evolution, and phenotypic impact of new genes. Genome Res. 20, 1313–1326.

47. Kalyaanamoorthy, S., Minh, B.Q., Wong, T.K.F., von Haeseler, A., and Jermiin, L.S. (2017). ModelFinder: fast model selection for accurate phylogenetic estimates. Nat. Methods 14, 587–589.

48. Katoh, K., and Standley, D.M. (2013). MAFFT multiple sequence alignment software version 7: improvements in performance and usability. Mol. Biol. Evol. 30, 772–780.

49. Khabbazian, M., Kriebel, R., Rohe, K., and Ané, C. (2016). Fast and accurate detection of evolutionary shifts in Ornstein-Uhlenbeck models. Methods Ecol. Evol. 7, 811–824.

50. Kleene, K.C. (2005). Sexual selection, genetic conflict, selfish genes, and the atypical patterns of gene expression in spermatogenic cells. Dev. Biol. 277, 16–26.

51. Kryuchkova-Mostacci, N., and Robinson-Rechavi, M. (2016). A benchmark of gene expression tissue-specificity metrics. Brief. Bioinform. 44, bbw008.

52. Kryuchkova-Mostacci, N., Robinson-Rechavi, M., Shmoish, M., Chalifa-Caspi, V., Shklar, M., and Ophir, R. (2016). Tissue-specificity of gene expression diverges slowly between orthologs, and rapidly between paralogs. PLOS Comput. Biol. 12, e1005274.

53. Kuleshov, M.V., Jones, M.R., Rouillard, A.D., Fernandez, N.F., Duan, Q., Wang, Z., Koplev, S., Jenkins, S.L., Jagodnik, K.M., Lachmann, A., et al. (2016). Enrichr: a comprehensive gene set enrichment analysis web server 2016 update. Nucleic Acids Res. 44, W90–W97.

54. Lan, X., and Pritchard, J.K. (2016). Coregulation of tandem duplicate genes slows evolution of subfunctionalization in mammals. Science 352, 1009–1013.

55. Langmead, B., and Salzberg, S.L. (2012). Fast gapped-read alignment with Bowtie 2. Nat. Methods 9, 357–359.

56. Leek, J.T., and Storey, J.D. (2007). Capturing heterogeneity in gene expression studies by surrogate variable analysis. PLoS Genet. 3, e161.

57. Leinonen, R., Sugawara, H., and Shumway, M. (2011). The Sequence Read Archive. Nucleic Acids Res. 39, D19–D21.

58. Liao, B.-Y., and Zhang, J. (2006). Low rates of expression profile divergence in highly expressed genes and tissue-specific genes during mammalian evolution. Mol. Biol. Evol. 23, 1119–1128.

59. Liu, X.-X., Zhang, H., Shen, X.-F., Liu, F.-J., Liu, J., and Wang, W.-J. (2015). Characteristics of testis-specific phosphoglycerate kinase 2 and its association with human sperm quality. Hum. Reprod. 126, dev301.

60. Marques, A.C., Dupanloup, I., Vinckenbosch, N., Reymond, A., and Kaessmann, H. (2005). Emergence of young human genes after a burst of retroposition in primates. PLoS Biol. 3, e357.

61. McCarrey, J.R., and Thomas, K. (1987). Human testis-specific PGK gene lacks introns and possesses characteristics of a processed gene. Nature 326, 501–505.

62. Mighell, A.J., Smith, N.R., Robinson, P.A., and Markham, A.F. (2000). Vertebrate pseudogenes. FEBS Lett. 468, 109–114.

63. Minh, B.Q., Nguyen, M.A.T., and von Haeseler, A. (2013). Ultrafast approximation for phylogenetic bootstrap. Mol. Biol. Evol. 30, 1188–1195.

64. Morel, B., Kozlov, A.M., Stamatakis, A., and Szöllősi, G.J. (2019). GeneRax: A tool for species tree-aware maximum likelihood based gene tree inference under gene duplication, transfer, and loss. BioRxiv 779066.

65. Necsulea, A., Soumillon, M., Warnefors, M., Liechti, A., Daish, T., Zeller, U., Baker, J.C., Grützner, F., and Kaessmann, H. (2014). The evolution of lncRNA repertoires and expression patterns in tetrapods. Nature 505, 635–640.

66. Nguyen, L.-T., Schmidt, H.A., von Haeseler, A., and Minh, B.Q. (2015). IQ-TREE: A fast and effective stochastic algorithm for estimating maximum-likelihood phylogenies. Mol. Biol. Evol. 32, 268–274.

67. Pace, J.K., Sen, S.K., Batzer, M.A., and Feschotte, C. (2009). Repair-mediated duplication by capture of proximal chromosomal DNA has shaped vertebrate genome evolution. PLoS Genet. 5, e1000469.

68. Podgornaia, A.I., and Laub, M.T. (2015). Pervasive degeneracy and epistasis in a protein-protein interface. Science 347, 673–677.

69. Pond, S.L.K., Frost, S.D.W., and Muse, S. V. (2005). HyPhy: hypothesis testing using phylogenies. Bioinformatics 21, 676–679.

70. Popescu, A.-A., Huber, K.T., and Paradis, E. (2012). ape 3.0: New tools for distance-based phylogenetics and evolutionary analysis in R. Bioinformatics 28, 1536–1537.

71. Potrzebowski, L., Vinckenbosch, N., Marques, A.C., Chalmel, F., Jégou, B., and Kaessmann, H. (2008). Chromosomal gene movements reflect the recent origin and biology of therian sex chromosomes. PLoS Biol. 6, e80.

72. Revell, L.J. (2012). phytools: an R package for phylogenetic comparative biology (and other things). Methods Ecol. Evol. 3, 217–223.

73. Rice, P., Longden, I., and Bleasby, A. (2000). EMBOSS: the European Molecular Biology Open Software Suite. Trends Genet. 16, 276–277.

74. Robinson, D.F., and Foulds, L.R. (1981). Comparison of phylogenetic trees. Math. Biosci. 53, 131–147.

75. Robinson, M.D., and Oshlack, A. (2010). A scaling normalization method for differential expression analysis of RNA-seq data. Genome Biol 11, 2010–2011.

76. Rogozin, I.B., Managadze, D., Shabalina, S.A., and Koonin, E. V (2014). Gene family level comparative analysis of gene expression in mammals validates the ortholog conjecture. Genome Biol. Evol. 6, 754–762.

77. Rohlfs, R. V., Harrigan, P., and Nielsen, R. (2014). Modeling Gene Expression Evolution with an Extended Ornstein–Uhlenbeck Process Accounting for Within-Species Variation. Mol. Biol. Evol. 31, 201–211.

78. Roy, S.W., and Gilbert, W. (2005). Rates of intron loss and gain: implications for early eukaryotic evolution. Proc. Natl. Acad. Sci. U. S. A. 102, 5773–5778.

79. Sanderson, M.J. (2002). Estimating absolute rates of molecular evolution and divergence times: a penalized likelihood approach. Mol. Biol. Evol. 19, 101–109.

80. She, X., Rohl, C.A., Castle, J.C., Kulkarni, A. V, Johnson, J.M., and Chen, R. (2009). Definition, conservation and epigenetics of housekeeping and tissue-enriched genes. BMC Genomics 10, 269.

81. Sikosek, T., Chan, H.S., and Bornberg-Bauer, E. (2012). Escape from Adaptive Conflict follows from weak functional trade-offs and mutational robustness. Proc. Natl. Acad. Sci. 109, 14888–14893.

82. Simão, F.A., Waterhouse, R.M., Ioannidis, P., Kriventseva, E.V., and Zdobnov, E.M. (2015). BUSCO: assessing genome assembly and annotation completeness with single-copy orthologs. Bioinformatics 31, 3210–3212.

83. Soubrier, J., Steel, M., Lee, M.S.Y., Der Sarkissian, C., Guindon, S., Ho, S.Y.W., and Cooper, A. (2012). The influence of rate heterogeneity among sites on the time dependence of molecular rates. Mol. Biol. Evol. 29, 3345–3358.

84. Starr, T.N., Picton, L.K., and Thornton, J.W. (2017). Alternative evolutionary histories in the sequence space of an ancient protein. Nature 549, 409–413.

85. Tria, F.D.K., Landan, G., and Dagan, T. (2017). Phylogenetic rooting using minimal ancestor deviation. Nat. Ecol. Evol. 1, 0193.

86. Van Raamsdonk, C.D., Bezrookove, V., Green, G., Bauer, J., Gaugler, L., O’Brien, J.M., Simpson, E.M., Barsh, G.S., and Bastian, B.C. (2009). Frequent somatic mutations of GNAQ in uveal melanoma and blue naevi. Nature 457, 599–602.

87. Veyrunes, F., Waters, P.D., Miethke, P., Rens, W., McMillan, D., Alsop, A.E., Grutzner, F., Deakin, J.E., Whittington, C.M., Schatzkamer, K., et al. (2008). Bird-like sex chromosomes of platypus imply recent origin of mammal sex chromosomes. Genome Res. 18, 965–973.

88. Warnefors, M., and Kaessmann, H. (2013). Evolution of the Correlation between Expression Divergence and Protein Divergence in Mammals. Genome Biol. Evol. 5, 1324–1335.

89. Wilson, B.A., Foy, S.G., Neme, R., and Masel, J. (2017). Young genes are highly disordered as predicted by the preadaptation hypothesis of de novo gene birth. Nat. Ecol. Evol. 1, 0146.

90. Yanai, I., Benjamin, H., Shmoish, M., Chalifa-Caspi, V., Shklar, M., Ophir, R., Bar-Even, A., Horn-Saban, S., Safran, M., Domany, E., et al. (2005). Genome-wide midrange transcription profiles reveal expression level relationships in human tissue specification. Bioinformatics 21, 650–659.

91. Yang, Z. (1994). Maximum likelihood phylogenetic estimation from DNA sequences with variable rates over sites: Approximate methods. J. Mol. Evol. 39, 306–314.

92. Yang, Z. (1995). A space-time process model for the evolution of DNA sequences. Genetics 139, 993–1005.

93. Yates, A., Akanni, W., Amode, M.R., Barrell, D., Billis, K., Carvalho-Silva, D., Cummins, C., Clapham, P., Fitzgerald, S., Gil, L., et al. (2016). Ensembl 2016. Nucleic Acids Res. 44, D710–D716.

94. Yu, G., Smith, D.K., Zhu, H., Guan, Y., and Lam, T.T.-Y. (2017). ggtree : an R package for visualization and annotation of phylogenetic trees with their covariates and other associated data. Methods Ecol. Evol. 8, 28–36.

95. Yu, Z., Morais, D., Ivanga, M., and Harrison, P.M. (2007). Analysis of the role of retrotransposition in gene evolution in vertebrates. BMC Bioinformatics 8, 308.

96. Zhang, L., and Li, W.-H. (2004). Mammalian housekeeping genes evolve more slowly than tissue-specific genes. Mol. Biol. Evol. 21, 236–239.

97. Zhang, Y.E., and Long, M. (2014). New genes contribute to genetic and phenotypic novelties in human evolution. Curr. Opin. Genet. Dev. 29, 90–96.

98. Zhang, Y.E., Landback, P., Vibranovski, M.D., Long, M., and Stark, A. (2011). Accelerated recruitment of new brain development genes into the human genome. PLoS Biol. 9, e1001179.

